# Developmental diversity and unique sensitivity to injury of lung endothelial subtypes during a period of rapid postnatal growth

**DOI:** 10.1101/2021.04.27.441649

**Authors:** Fabio Zanini, Xibing Che, Carsten Knutsen, Min Liu, Nina E. Suresh, Racquel Domingo-Gonzalez, Steve H. Dou, Daoqin Zhang, Gloria S. Pryhuber, Robert C. Jones, Stephen R. Quake, David N. Cornfield, Cristina M. Alvira

**Author notes:** Senior author (Stephen R. Quake, David N. Cornfield, Cristina M. Alvira). Correspondence, Address for Correspondence: Cristina M. Alvira, M.D., Stanford University School of Medicine, Center for Excellence in Pulmonary Biology, 770 Welch Road, Suite 435, Palo Alto, CA 94304.

## Abstract

At birth, the lung is still immature, heightening susceptibility to injury but enhancing regenerative capacity. Angiogenesis drives postnatal lung development. Therefore, we profiled the transcriptional ontogeny and sensitivity to injury of pulmonary endothelial cells (EC) during early postnatal life. Although subtype speciation was evident at birth, immature lung EC exhibited transcriptomes distinct from mature counterparts, which progressed dynamically over time. Gradual, temporal changes in aerocyte capillary EC (CAP2), contrasted with more marked alterations in general capillary EC (CAP1) phenotype, including distinct CAP1 present only in the early alveolar lung expressing *Peg3*, a paternally imprinted transcription factor. Hyperoxia, an injury which impairs angiogenesis, induced both common and unique endothelial gene signatures, dysregulated capillary EC cross-talk, and suppressed CAP1 proliferation while stimulating venous EC proliferation. These data highlight the diversity, transcriptomic evolution, and pleiotropic responses to injury of immature lung EC, possessing broad implications for lung development and injury across the lifespan.

## Introduction

At birth, rhythmic distention of the lungs and a marked rise in oxygen tension rapidly decrease pulmonary vascular resistance to increase pulmonary blood flow. Lining the pulmonary blood vessels, endothelial cells (EC) serve as first responders to these dramatic changes in oxygen and shear stress by increasing the production of endothelial-derived vasodilators and altering their cell shape to facilitate successful transition from the fetal to neonatal circulation^1^. Subsequently, rapid expansion of the pulmonary microcirculation during early postnatal life drives the formation of hundreds of millions of alveoli to dramatically increase gas exchange surface area^2^.

Completion of parenchymal and vascular development after birth renders the immature lung highly susceptible to environmental insults that can disrupt lung development. This is frequently observed in the context of preterm birth, where disrupted pulmonary angiogenesis and alveolarization results in the chronic lung disease, bronchopulmonary dysplasia (BPD), the most common complication of prematurity^3^. However, even in the mature lung, developmental mechanisms and gene expression programs can be reactivated to promote lung regeneration after injury^4, 5^. Thus, a detailed understanding of pulmonary EC diversity and phenotypic changes across postnatal development has important implications for lung diseases affecting not only premature infants, but also older infants, children, and adults.

EC are both highly heterogeneous, often displaying multiple phenotypes even within a single organ, and tremendously plastic in response to changes in the microenvironment. Single cell transcriptomics has provided a high-resolution picture of the adult lung in mice^6^ and humans^7^ and begun to shed light on specific populations in the epithelial^8^, immune^9^, mesenchymal^10^ and endothelial compartments^9, 11, 12^. However, deep interrogation of EC phenotypic transitions during this critical window of postnatal development, and the unique responses of specific endothelial subtypes to injury has not been previously reported. In this study we utilized plate-based, single cell RNA-sequencing (scRNA-seq), fluorescent *in situ* hybridization (FISH) in murine and human lung tissue, and mechanistic studies in primary lung EC to identify developmental alterations in the EC transcriptome during early postnatal life and to assess how EC diversity and the phenotype of select EC subtypes are altered by hyperoxia, an injury that disrupts both lung parenchymal and vascular growth.

## Results

### Pulmonary endothelial heterogeneity increases after birth

To define phenotypic changes of pulmonary endothelial cells across early postnatal development, we isolated lungs from C57BL/6 mice at E18.5 (early saccular), P1 (late saccular), P7 (early alveolarization) and P21 (late alveolarization), and P7 mice exposed to chronic hyperoxia, an injury that impairs alveolarization and angiogenesis (Figure 1A). After optimizing lung dissociation to maximize EC viability and yield at each timepoint^9^, we perfused the pulmonary circulation, digested the lungs and performed fluorescent activated cell sorting (FACS) to select CD31+ EC^9^. We transcriptionally profiled 2931 EC from 10 C57BL/6 mouse lungs using Smart-Seq2 at a depth of approximately 1 million reads per cell. These methodologies resulted in detection of 4376 genes per cell (median), more than twice the number of genes per EC detected in recent single cell reports from the adult^6, 13^ and developing lung^11, 14, 15^ (Figure 1B).

**Figure 1.**
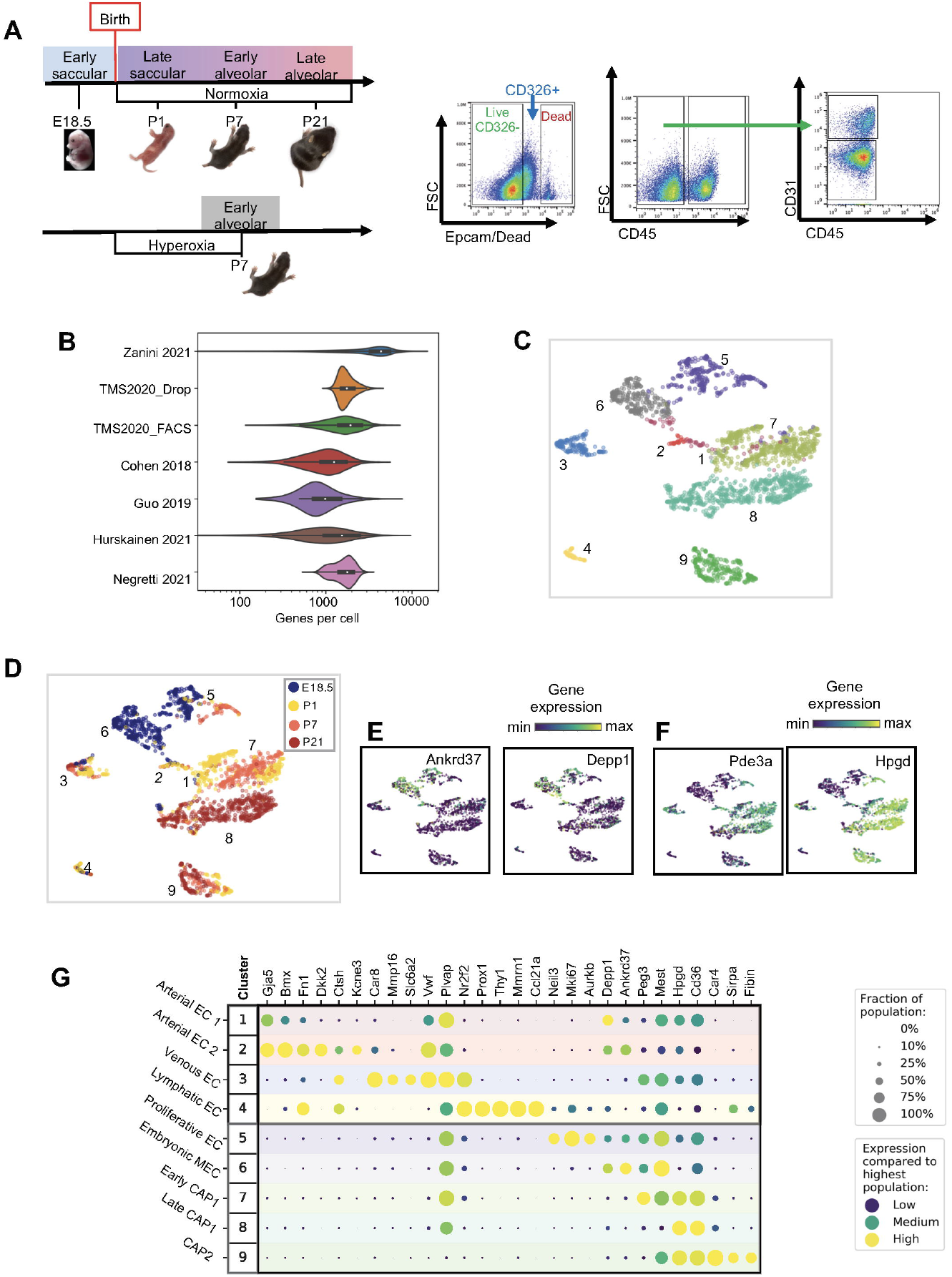
Pulmonary endothelial heterogeneity increases after birth. (A) Overview of the experimental design including the four timepoints (E18.5, P1, P7, P21) corresponding to key stages in late lung development. In separate experiments, pups were maintained in chronic hyperoxia from birth until P7. Lungs were isolated, perfused, and digested and endothelial cells isolated by fluorescence activated cell sorting (FACS) for the live, CD326^-^CD45^-^CD31^+^ population. (B) Violin plots displaying the number of genes per detected per cell in our data set compared to recently published reports. (C) UMAP and Leiden clustering of 2931 endothelial cells identifying 9 transcriptionally distinct EC subpopulations present during the lung from the early saccular to late alveolar stages of development. (D) UMAP of EC clusters identifying developmental timepoint of cell origin with E18.5 (blue), P1 (yellow), P7 (orange) and P21 (red). (F) UMAP plots of genes expressed predominantly before (E) and (F) after birth. For (E) and (F), the color scale is a logarithmic scale with a pseudocount of 0.1 counts per million, normalized to the highest expressing cell. (G) Dot plot showing level of expression (purple to yellow), and fraction of each population expressing the particular gene (dot size) for distinguishing genes expressed by each cluster. See also Figure S1.

We visualized the data with Uniform Manifold Approximation and Projection (UMAP)^16^, and performed unsupervised clustering using the Leiden algorithm^17^ to reveal nine EC subtypes in the perinatal lung (Figure 1C), including four clusters of macrovascular EC, one cluster of proliferative EC, and four microvascular EC (MEC) clusters. Visualization of individual cells by developmental stage identified significant changes in lung heterogeneity and shifts in the EC landscape during this transition from late embryonic to early postnatal life (Figure 1D). These included the disappearance of one cluster present only at E18.5, (Cluster 6), the appearance of three postnatal microvascular EC clusters comprised primarily of cells from specific developmental timepoints (Clusters 5, 6, 7 and 8), and a fourth microvascular cluster exhibiting a more gradual developmental gradient (Cluster 9). These developmental shifts were accompanied by significant gene expression changes, including genes primarily expressed prior to birth (e.g. *Ankrd37* and *Depp1*) (Figure 1E), versus others expressed only after birth (e.g. *Pde3a* and *Hpgd*) (Figure 1F).

We were able to annotate these nine EC clusters by comparing distinguishing marker genes in each group. Arterial EC (Clusters 1 and 2) shared expression of *Gja5*, a gene that modulates arterial identity^18^ and *Bmx*, a tyrosine kinase expressed in large arteries^19^, but were split by the clustering algorithm into two subtypes (Art 1 and 2) distinguished by the expression of the potassium channel *Kcne3*, and *Dkk2*, an inhibitor of canonical Wnt signaling (Figure 1G). Venous EC (Cluster 3) shared expression of *Vwf* with arterial EC, but were distinguished by the expression of *Nr2f2*, a transcription factor (TF) that suppresses Notch signaling to confer venous identity^20^. Venous EC also uniquely expressed the norepinephrine transporter, *Slc6a2*, and high levels of *Mmp16*, and *Car8*. Lymphatic EC (Cluster 4) were identified by the exclusive expression of canonical genes *Thy1*^21^ and *Prox1*^22^ in addition to *Ccl21a*. Proliferating EC (Cluster 5) were distinguished by high expression of *Mki67* and numerous additional genes associated with mitosis, with the cluster comprised primarily of microvascular EC, and a smaller number of macrovascular EC (Figure S1A). Proliferating EC exhibited two peaks in abundance, one occurring just prior to birth, and the second at P7 (Figure S1B and C). Consistent with recent reports, the embryonic microvascular EC (Cluster 6) separated into two coarse cell populations after birth, one (Cluster 9) distinguished by high expression of carbonic anhydrase 4 (*Car4*), consistent with aerocyte (CAP2) identity, and two separate clusters (7 and 8) expressing *Plvap*, consistent with general capillary (CAP1) identity^23^.

### Pulmonary EC in the perinatal lung are transcriptionally distinct from the adult

We next determined how similar perinatal lung EC are to adult lung EC by directly comparing our data with the cell atlas from adult mice, Tabula Muris (TM)^6^ (Figure 2A). Harmonization of the two data sets highlighted the marked difference between immature lung EC compared to their mature counterparts, particularly within the microvasculature. Perinatal lymphatic EC embedded close to adult lymphatic EC, suggesting a similar phenotype (Figure 2B). Perinatal arterial EC also embedded close to adult arterial EC, with our Art II subtype appearing most similar to the adult. Interestingly, perinatal venous EC did not embed near adult venous EC, suggesting significant alterations in venous phenotype during postnatal development. Although the perinatal CAP2 also embedded close to adult CAP2, a developmental gradient was evident (Figure 2C). This developmental progression was more striking in the CAP1 EC, which formed separate, adjacent clusters that layered developmentally, beginning with the embryonic EC and ending with late CAP1 embedding closest to, but still separate from adult CAP1 EC (Fig 2B and C). Proliferative EC were an entirely unique cluster in our data set, confirming the absence of proliferation in the healthy adult pulmonary endothelium.

**Figure 2.**
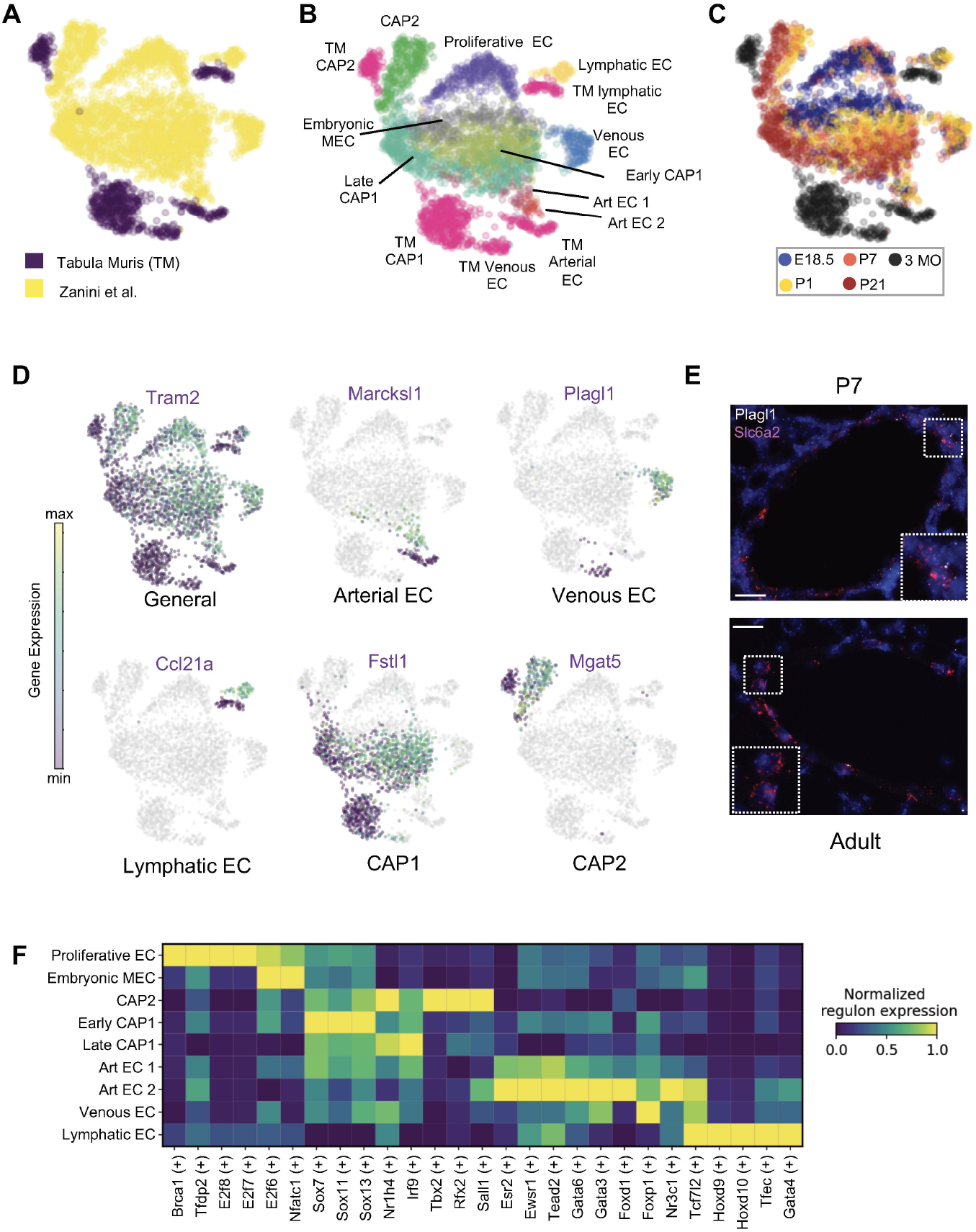
Pulmonary EC in the perinatal lung are transcriptionally distinct from the adult. Our data set and lung data from Tabula Muris (TM) were merged, restricted to genes that were overlapping, normalized to counts per million reads, feature selected and subjected to principal component analysis (PCA), followed by UMAP. UMAP plots of (A) data source (B) cell type and (C) developmental stage using our data combined with Tabula Muris. (D) UMAP plots of select genes differentially expressed in perinatal EC clusters versus analogous clusters in TM, with gene expression colored logarithmically as described above. (E) Multiplexed *in situ* hybridization to detect expression of the venous EC marker *Slc6a2* (red), and *Plagl1* (white) in lung tissue from mice at P7 and adult, with high magnification insets outlined with the dotted square. Cal bar=20μm. (F) Heat map of select TF regulons enriched in each EC subtype as identified by SCENIC, normalized such that the EC subtype exhibiting the highest enrichment for a specific regulon is given a score of 1.

We next identified genes differentially expressed by the perinatal and adult EC subtypes. One group of genes appeared to be generally up-regulated in perinatal lung EC as compared to the adult, including *Tram2*, a gene required for type I collagen synthesis^24^, a matrix component produced in large quantities during lung development^25^ (Figure 2D). In contrast, other genes were specifically up-regulated in select perinatal compared to adult EC subtypes. For example, *Marckls1*, a regulator of developing blood vessel diameter, was more highly expressed in perinatal compared to adult arterial EC^26^. In perinatal venous EC, *Plagl1*, a zinc finger TF that regulates proliferation, cell adhesion and extracellular matrix composition^27^, was up-regulated. Perinatal lymphatic EC exhibited high expression of the lymphocyte/dendritic cell chemoattractant *Ccl21a*^28^, perhaps contributing to the gradual increase in resident lung lymphocytes and dendritic cells observed across the first three weeks of postnatal life^9^. The perinatal MEC also expressed genes unique from adult MEC, including *Fstl1* by perinatal CAP1, encoding a secreted ECM glycoprotein that promotes EC survival, migration and differentiation^29^, and *Mgat5* by perinatal CAP2, a gene that enhances VEGFR2 signaling^30^, a pathway essential for CAP2 survival^31^.

To confirm that these developmental differences in gene expression represented true biologic findings and not batch effects, we validated these results via FISH to detect the combination of *Plagl1* with the venous EC specific marker *Slc6a2* (Figure 1D) in pulmonary veins of P7 and adult mice. *Slc6a2* expression was apparent in venous EC from both P7 and adult mice (Figure 2E). However, although *Plagl1* expression was broadly detected in venous EC from P7 mice, it was virtually absent in adult venous EC (Figure 2E).

We next performed gene regulatory network analysis using SCENIC^32^ to identify putative transcription factors (TFs) fingerprints uniquely enriched in each perinatal EC subtype. This method allowed the visualization of both shared and unique regulatory networks (regulons) in similar and distinct EC subtypes (Figure 2F). For example, the proliferative EC exhibited gene enrichment in regulons with established roles in cell proliferation including *Brca1*^33^ and multiple members of the E2F family (e.g. *E2f6, E2f7*, and *E2f8*)^34^. Of note, the embryonic MEC shared some regulon enrichment with proliferative EC, including *E2f6*, and *Nfatc1*, a TF required for vascular patterning and VEGF-induced angiogenesis^35^. The postnatal MEC also shared enrichment with a cluster of regulons (e.g. *Sox7, Sox11, Sox3, Nr1h4*, and *Irf9*). Further, the CAP2 were enriched in regulons (e.g. *Tbx2, Rfx2*, and *Sall1*) not shared with other EC. The macrovascular EC generally exhibited more heterogeneity in predicted TF networks but shared enrichment of *Tcf7l2*, an effector of canonical Wnt signaling, perhaps serving to promote vascular stability in more established vessels^36^. The Art2 were specifically enriched in regulons directed by *Gata6*, a TF that promotes EC angiogenic function^37^, and *Gata3*, a mediator of Tie2 expression in macrovascular EC^38^. Venous EC shared enrichment of the *Gata3* regulon, albeit at a lower level, but unique enrichment of the regulon of *Foxp1*, a TF that limits vascular inflammation in response to shear stress^39^. Lymphatic EC were enriched for pathways known to influence lymphatic development and function including *Hoxd10*, a TF that regulates responses to VEGF-C^40^. Importantly, Art1 EC exhibited shared enrichment of both micro- and macrovascular regulons, suggesting an intermediate phenotype. Taken together, these data highlight the marked heterogeneity of the perinatal lung EC, and demonstrate that the majority of perinatal EC subtypes are transcriptionally distinct from, but converging towards their adult counterparts slowly across postnatal development.

### Arterial EC exhibiting a distinct transcriptome are present in the late embryonic and early postnatal lung

The presence of a transcriptionally distinct subtype of arterial EC has not been previously described. In contrast to relatively stable abundances of the lymphatic, venous and Art2 EC, the Art1 EC almost completed disappeared between P7 and P21 (Figure 3A). To validate our computational findings and gain greater clarity on the function of these cells, we localized Art 1and 2 *in situ* in the P1 lung by simultaneously detecting *Cdh5, Gja5*, and *Dkk2*, and identified arterial EC lining larger arteries and small arterioles as exclusively Art2. In contrast, single positive Art1 EC expressing *Gja5* but not *Dkk2* were located in the distal lung parenchyma, adjacent to *Gja5*^-^*Cdh5*^+^ microvascular EC (Figure 3B). Similar results were observed when employing another Art2 marker, *Mgp* in combination with *Gja5* (See Figure S2A). Of note, we performed similar studies at P7, and found that at this later timepoint, the *Gja5*+ EC in the distal lung parenchyma also expressed *Dkk2* (See Figure S2B). Given this inconsistency with our abundance data, we looked more closely at the original embedding and noted that the clustering algorithm identified a number of P7 cells as Art1 that actually embedded with the early CAP1 and expressed many CAP1 genes, suggesting that their biologic identity is actually CAP1. We also re-analyzed a separate single cell transcriptomic dataset from the neonatal lung that contained cells from P3, and found that in this dataset that the arterial cluster clearly contained a similar mixed population, with a clear subpopulation of *Gja5*+ EC that lacked expression of *Dkk2* and *Kcne3*, consistent with the Art1 phenotype (See Figure S2C)^15^.

**Figure 3.**
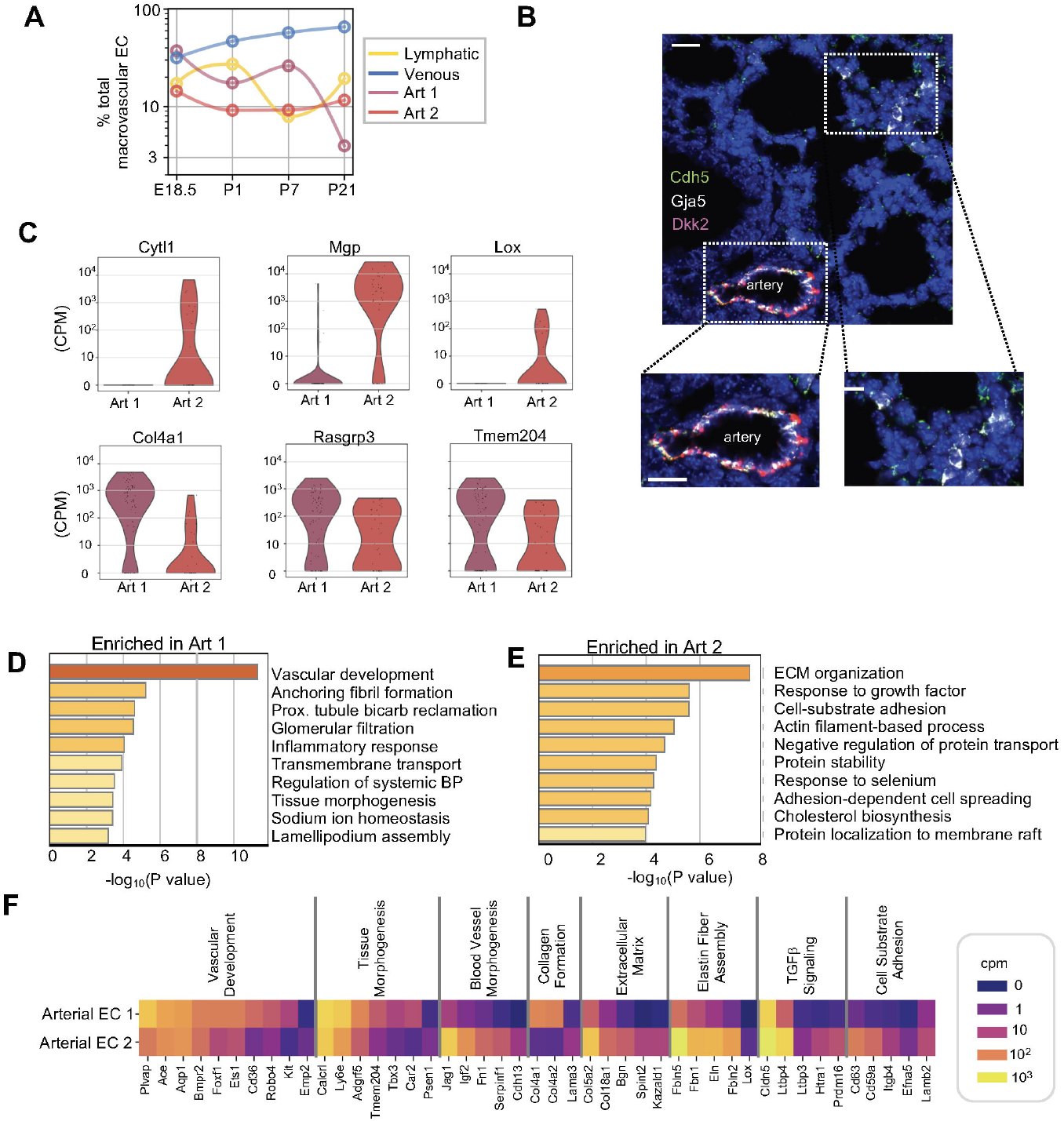
Arterial EC exhibiting a distinct transcriptome are present in the late embryonic and early postnatal lung. (A) Quantification of the abundance of each macrovascular EC subset at each developmental timepoint expressed on a logarithmic scale as percentage of total macrovascular EC. (B) Multiplexed *in situ* hybridization to detect expression of *Cdh5* (green), *Gja5* (white) and *Dkk2* (red) in lung tissue from mice at P1. Cal bar=20μm. (C) Violin plots depicting the gene expression of genes highly expressed by Arterial II (red) versus Arterial I (purple) EC. (D) Pathway analysis performed via Metscape on the top 100 up-regulated genes in the Art l and (E) Art II clusters, respectively. (F) Heatmap of select differentially expressed genes in Art I and II identified within enriched pathways illustrated. The color scale is logarithmic counts per million, normalized to the highest expressing population for each gene. See also Figure S2.

In addition to the differences in *Dkk2* and *Kcne3* expression, Art1 also lacked expression of additional genes highly expressed by Art2 cells including *Cytl1*, recently identified as a large artery marker gene ^41^, *Mgp*, a BMP antagonist^42^, and *Lox*, an enzyme required for elastogenesis ^43^ (Figure 3C). In contrast, Art1 EC highly expressed *Col4a1, Rasgrp3*, a guanine exchange factor downstream of VEGF signaling ^44^, and *Tmem204*, a hypoxia regulated transmembrane protein that regulates VEGF signaling^45^.

Differentially expressed genes in Art1 and 2 EC were enriched for distinct pathways related to vascular growth and remodeling (Figure 3D-F and Table S1). In Art1 EC, the highest enrichment was found in “vascular development”, including high expression of *Col4a1* and *Col4a2*, integral components of the vascular basement membrane ^46^, angiogenic genes such as *Ets1*^47^, *Robo4*^48^, *Foxf1*^49^, and *Kit*^50^, and *Igfbp7*, a secreted angiocrine factor that may play a role in vascular patterning by preventing excess VEGF-A bioavailability^51^. Extracellular matrix organization and elastin fiber assembly were enriched in Art II EC, including high expression of *Bgn*, an ECM component that enhances HIF-1α-mediated transcription of VEGF^52^, and multiple additional genes important for elastin fiber assembly including *Eln* and *Fbln2*^53^.

### CAP1 derived from the early alveolar lung are marked by high expression of paternally expressed gene-3 (Peg3)

We then focused our attention on the microvascular EC populations, three of which were associated with specific timepoints. Embryonic MEC represented almost 60% of total EC just prior to birth (Figure 4A), and were distinguished from postnatal MEC (clusters 7-9) by the high expression of *Mest*, a paternally imprinted gene, *Depp1*, a hypoxia-induced gene that regulates autophagy^54^, *Sparcl1*, a matricellular protein that enhances mural cell recruitment^55^, and the HIF-1α target *Rgcc*, a complement response gene that promotes NFκB-mediated VEGFR2 expression^56^ (Figure 4B). Abundance of embryonic MEC dropped dramatically after birth (Figure 4A), replaced over the first week of life by increasing abundance of CAP1 and CAP2 EC. Although the aCAP were present at all postnatal timepoints (see below), early CAP1 (Cluster 7) consisted almost exclusively of EC from P1 and P7 (Figure 1D). This subtype itself was eventually replaced by late CAP1 (Cluster 8), almost exclusively comprised of EC from P21. We performed RNA velocity^57^ on both CAP1 subtypes and reconstructed a cellular trajectory from early to late CAP1 along the embedding, with a general downwards flow, suggestive of continuous progression (Figure 4C). Taken together these data suggest that these two transcriptional clusters represent different developmental states of the CAP1. Further analysis identified an overall decrease in the number of genes expressed by late versus early CAP1 (Figure 4D), consistent with a reduction in transcription as CAP1 progress towards quiescence.

**Figure 4.**
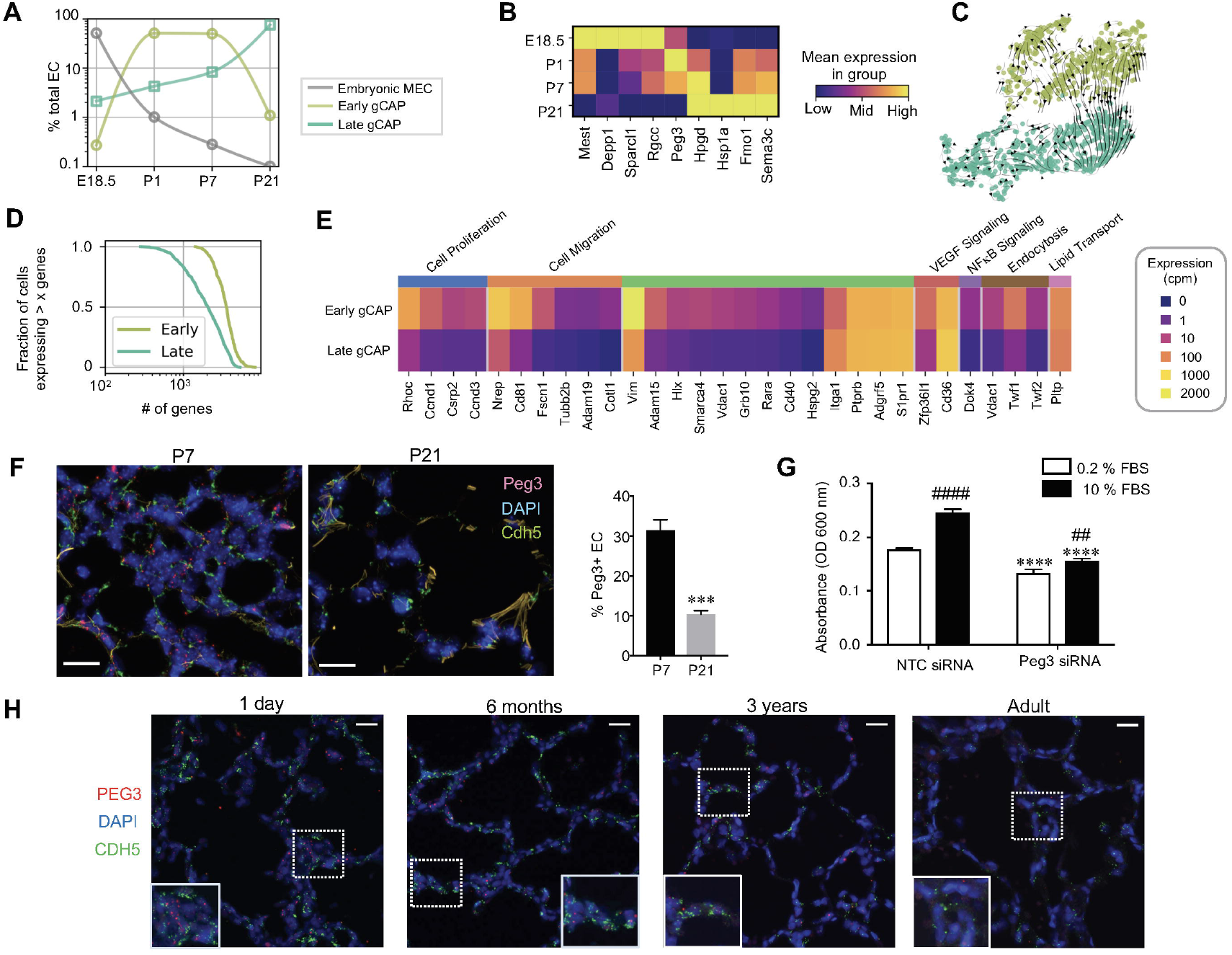
CAP1 derived from the early alveolar lung are marked by high expression of paternally expressed gene-3 (Peg3). (A) Relative abundance of embryonic, early CAP1, and late CAP1 at each developmental timepoint expressed on a logarithmic scale as percentage of total EC. (B) Heatmap of genes highly expressed in embryonic MEC, early and late CAP1 across the four timepoints, where the bottom and the top of the color scale are set to the geometric mean expression of the lowest and highest expressing timepoint, respectively. (C) RNA velocity demonstrating a transition from early to late CAP1. (D) Cumulative distribution of number of expressed genes, indicating early CAP1 express more genes than late CAP1. (E) Heatmap of genes differentially expressed between early and late CAP1, grouped by select pathways. (F) Multiplex *in situ* hybridization (middle) to detect *Peg3* (red) and *Cdh5* (green) in lung tissue at P7 and P21. Calibration bar=20μm. Quantification (right) of the percent of *Peg3+* EC versus total EC in lung sections from P7 and P21 mice (31.4±1.6 vs. 10.3±0.55 and P=0.002 by t-test with n=3 mice per group). (G) Proliferation of primary EC transfected with NTC or Peg3 siRNA and stimulated with either 0.2% or 10% FBS, with ****p<0.0001 versus NTC transfected cells, and ##p<0.01 and ####p<0.0001 vs. 0.2% FBS via 2-Way ANOVA. Data is representative of 3 independent experiments, with 8 technical replicates per condition. (H) Representative images of multiplex *in situ* hybridization to detect *PEG3* (red) and *CDH5* (green) RNA in human lung tissue from donors of various ages extending from early alveolarization to early adulthood with high magnification insets of lung MEC outlined by the dotted squares. Cal bar=20μm. See also Figure S3-S5.

Pathway analysis of differentially expressed genes demonstrated that early CAP1 exhibited enrichment in pathways regulating actin cytoskeletal organization, including *Cotl1*, a gene that promotes lamellipodia protrusion^58^, and multiple genes that regulate cell motility including *Fscn1, Twf1, Twf2, Cd81* and *Nrep* (Figure 4E and Figure S3). Genes that promote proliferation were also increased including *Ccnd1, Ccnd3, and Csrp2*, as were multiple angiogenic genes including *Rara*, the gene encoding the retinoic acid receptor alpha^59^, *Adam15*, a metalloproteinase that promotes EC survival^60^ and retinal neovascularization^61^, and *Cd40*, a cell surface receptor that enhances VEGF expression^62^. Late CAP1 were characterized by a general down-regulation of these genes, in concert with a significant up-regulation of *Cd36*, a receptor for thrombospondin that inhibits EC migration^63^, *Itga1*, a component of the α1β1 integrin that binds the basement membrane components collagen IV and laminin^64^, and *S1pr1*, encoding a G-protein coupled receptor that enhances EC barrier function and stability of nascent vessels^65^. Consistent with a recent report identifying a subset of gCAP that possess progenitor properties and co-express *Foxf1* and *Kit*^66^, we determine the fraction of *Foxf1*+*Kit*+ EC in each cluster, and found that the early CAP1 contained the highest fraction of these cells (Figure S4).

Further analysis of genes differentially expressed between the early and late CAP1 identified the paternally expressed gene-3 (*Peg3*) as one of the most highly expressed genes by early CAP1 (Figure 1D and 4B). *Peg3* is an imprinted gene that encodes a zinc finger protein highly expressed during embryogenesis^67^ that marks cells capable of self-renewal in adult mouse tissue^68^. We employed *in situ* hybridization to detect *Peg3* expression in CAP1 from mice at P7 and P21. In agreement with our computational data, *Peg3*+ CAP1 were abundant throughout the distal pulmonary endothelium at P7. In contrast, *Peg3*+ CAP1 were rare in the P21 lung, with 3-fold greater *Peg3*+ CAP1 present at P7 versus P21 (Figure 4F). Although it has been previously shown that high *Peg3* expression is observed in EC with high proliferative potential, the specific role for *Peg3* in driving proliferation in pulmonary EC in the developing lung has not been studied. Therefore, to further clarify the function of *Peg3* in the early CAP1, we isolated primary microvascular EC from P7 mice as we have described previously^69–72^. Immunostaining these primary EC to detect PEG3 protein identified numerous cells expressing high levels of nuclear PEG3 in culture, that was effectively diminished by transfecting the cells with *Peg3* siRNA (Figure S5). Silencing *Peg3* blunted basal proliferation, and significantly diminished proliferation stimulated by 10% FBS (Figure 4G).

Remarkably, the developmental expression of *Peg3* in microvascular EC was conserved in the human lung. At 1 day of life, the majority of EC in the distal lung co-expressed *PEG3*. *PEG3* expression remained moderately high in the human lung at 6 months of age, a timepoint corresponding to ongoing alveolarization and pulmonary angiogenesis. However, *PEG3* expression was reduced in the lung EC by 3 years of age, a time corresponding to the end of bulk alveolarization, and minimally expressed in human lung EC at early adulthood (Figure 4H).

### CAP2 EC exhibit dynamic developmental alterations in gene expression to influence cell migration, angiogenesis and immunity

Recent reports have identified CAP2 as specialized MEC that require alveolar epithelial type I-derived VEGF for survival^31^, and appear to play a key role in both alveolarization and lung repair after viral infection^73^. However, the specific functions of CAP2 during lung development have not been identified. Although the CAP2 cluster contained EC derived from all three postnatal timepoints, a clear developmental gradient was apparent, confirmed by both pseudotime analysis and RNA velocity (Figure 5A). Identification of genes that were highly expressed at each of the three postnatal timepoints included high expression of *Gap43*, and *Nid2* at P, *Tspan12, Hpgd* and *Cav2* at P7, and *Fibin* and *Car4* at P21 (Figure 5B). Identification of select genes exhibiting varied expression over pseudotime demonstrated a pattern of a small number of highly expressed genes slowly increasing, but a more dramatic decrease in a greater number of genes (See Figure S6).

**Figure 5.**
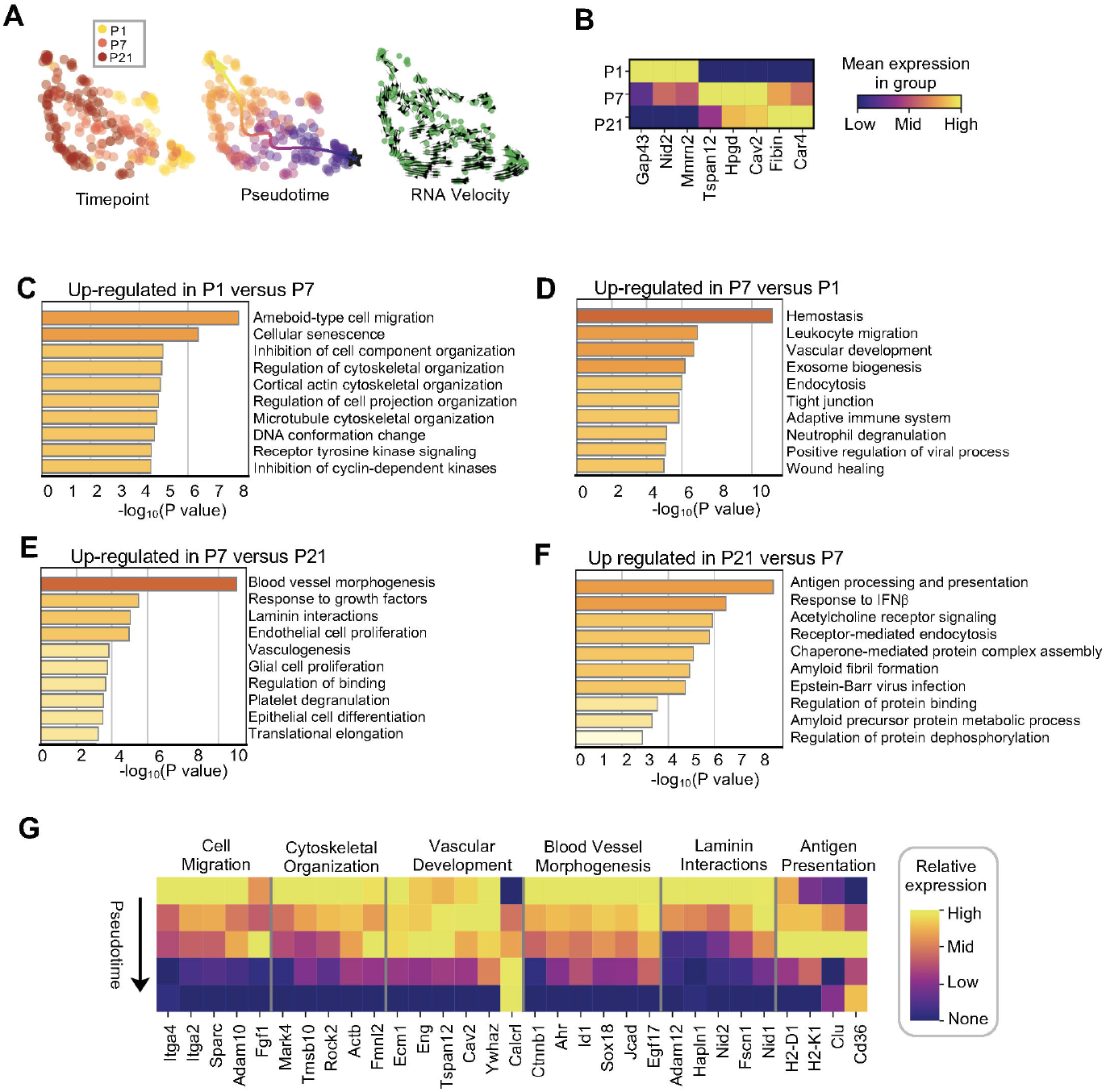
CAP2 exhibit dynamic developmental alterations in gene expression to influence cell migration, angiogenesis and immunity. (A) UMAP plots of genes highly expressed by CAP2, with gene expression colored logarithmically as described above. (B) Heatmap of genes highly expressed by CAP2 at each postnatal timepoint, where the bottom and the top of the color scale are set to the geometric mean expression of the lowest and highest expressing timepoint, respectively. (C) UMAP embedding of CAP2 colored by timepoint (left), by pseudotime starting from the bottom-right corner with an average trajectory (middle), and with all EC in green with RNA velocity overlayed (right). All three methods indicate maturation along the developmental axis. (C-F) Pathway analysis of DEG in the CAP2 between each of the postnatal timepoints. (G) Heatmap of select DEG in CAP2 across pseudotime within enriched pathways. See also Figure S6.

Further interrogation of phenotypic changes across early postnatal development were inferred by pathway analysis of the DEGs at each developmental timepoint (Figures 5C-F) and the visualization of select genes that changed across pseudotime (Figure 5G). These studies revealed enriched expression of genes important for cell migration and cytoskeletal remodeling, including *Adam10*, a metalloproteinase that promotes EC permeability^74^, and *Itga2* and *Itga4*, components of the integrin receptors for collagen and fibronectin at P1. By P7, genes related to vascular development were upregulated, including *Eng*, an ancillary TGFβ receptor essential for extraembryonic angiogenesis^75^, *Tspan12*, a tetraspanin that regulates retinal vascular development by augmenting Norrin signaling^76^, and *Ywhaz*, encoding a 14-3-3 adapter protein required for lung vascular development^77^. By P21, these, and other genes important for blood vessel morphogenesis were down-regulated, while genes regulating antigen processing and presentation were increased, including up-regulation of *Cd36* as seen in CAP1, a gene that functions as a scavenger receptor involved in the presentation of MHC class II antigens^78^, and *H2-D1* and *H2-K1*, genes encoding MHC Class I molecules.

### Chronic hyperoxia delays microvascular maturation, induces both common and unique EC gene signatures, and disrupts crosstalk between CAP1 and CAP2 EC

We then performed single cell transcriptomics on pulmonary EC isolated from pups exposed to chronic hyperoxia during the first seven days of life (Figure 1A), an injury that disrupts postnatal parenchymal and vascular development^69, 79, 80^. Even at this early timepoint, hyperoxia grossly diminished the complexity of the pulmonary circulation apparent after latex dye perfusion (Figure S7). Hyperoxia altered both the relative abundance and gene expression within select EC subtypes. Dimensionality reduction via UMAP including only EC from P7 revealed well mixed macrovascular EC but greater segregation within microvascular EC (Figures 6A and B). Early CAP1 were slightly increased, while late CAP1 were less abundant (Figure 6C). In addition, a novel cluster of proliferating venous EC was present, comprised almost exclusively by EC from hyperoxia-exposed mice.

**Figure 6:**
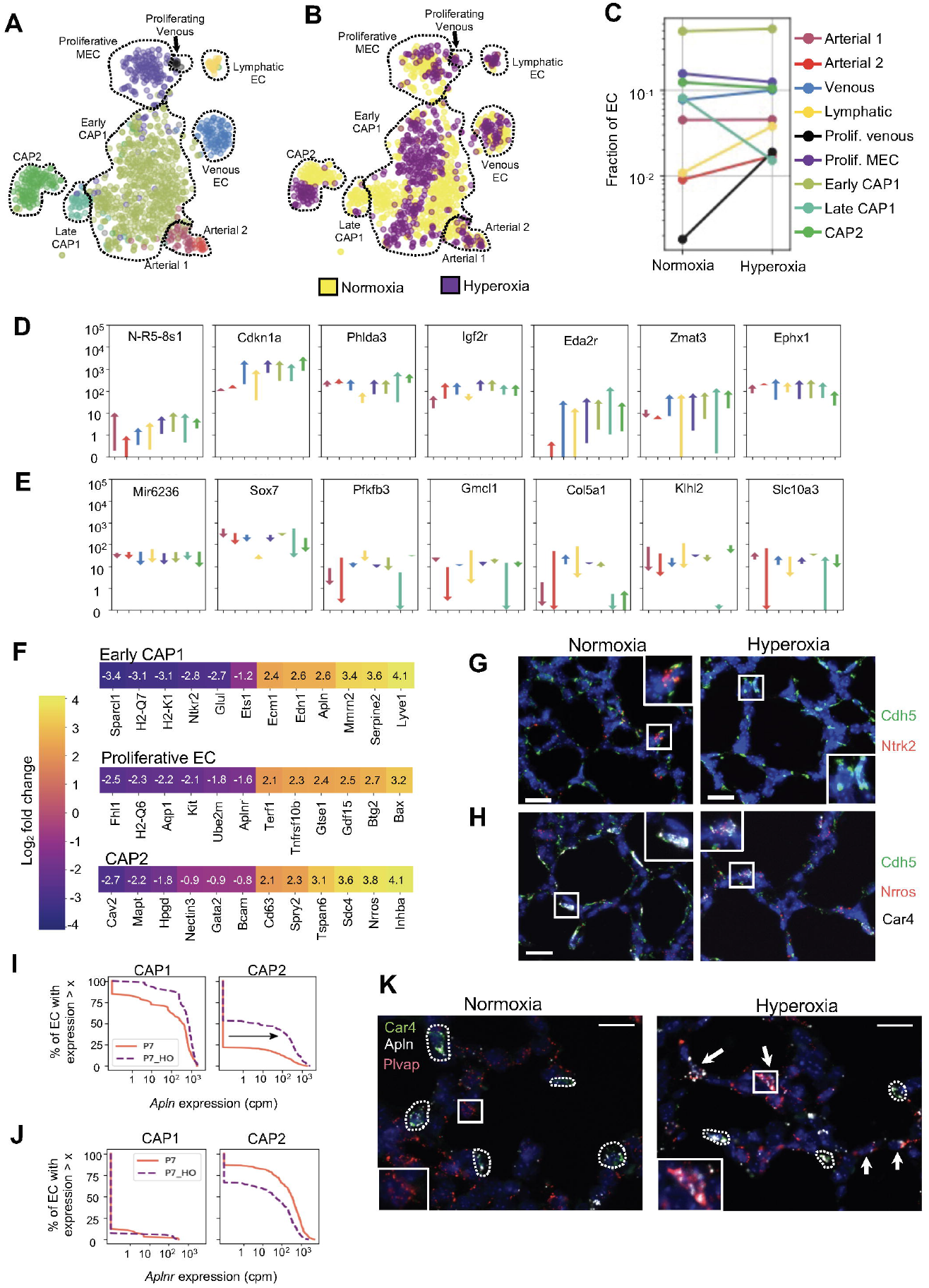
Chronic hyperoxia delays microvascular maturation, induces both common and unique EC gene signatures, and disrupts crosstalk between CAP1 and CAP2 EC. (A) UMAP and unsupervised clustering of EC derived from the P7 lung maintained in normoxia and hyperoxia defining 9 EC clusters. (B) UMAP plot colored by treatment condition with normoxia in yellow and hyperoxia in purple. (C) Relative abundance of each EC subtype as a fraction of total EC in normoxia versus hyperoxia, shown on a logarithmic scale. (D-E) Identification of common up- (D) and down- (E) regulated genes in hyperoxia. In each plot, the start of the arrow corresponds to the average expression of the gene in normoxia, and the tip of the arrow corresponds to the average expression of the gene in that cluster in hyperoxia. The y axis uses a logarithmic scale of gene expression in cpm, with the arrow colored to match each individual cluster. (F) Heat map of select up- and down-regulated genes in hyperoxia versus normoxia at P7 in the early CAP1 (top), proliferative EC (middle), and CAP2 (bottom). Values in each square represent the log2 fold changes in average expression. (G) Representative multiplex *in situ* hybridization to detect *Cdh5* (green) and *Ntrk2* (red) in combination with DAPI (blue) in lung tissue from P7 mice exposed to normoxia or hyperoxia. Calibration bar=20μm. (H) Representative multiplex in situ hybridization to detect *Cdh5* (green) *Nrros* (red), and *Car4* (white) in combination with DAPI (blue) in lung tissue from P7 mice exposed to normoxia or hyperoxia. Calibration bar=20μm. (I-J) Cumulative distributions of expression of *Apln* (I) and *Aplnr* (J) in CAP2 and early CAP1 at P7 in normoxia and hyperoxia. The black arrow highlights the proportion of early CAP1 expressing *Apln* in hyperoxia. (K) Representative multiplex *in situ* hybridization to detect *Car4* (green), *Apln* (white), and *Plvap* (red) in lung tissue from P7 mice exposed to normoxia or hyperoxia. Cal bar=20μm. See also Figure S7 and S8.

Despite the transcriptional heterogeneity of lung EC present at P7, hyperoxia induced a common gene signature across all EC subtypes (Figure 6D and E). Commonly up-regulated genes included *Cdkn1a*, an inhibitor of cell proliferation that also limits oxidative stress^81^, *Eda2r*, a member of the TNF receptor superfamily that is increased in the aging lung^82^, and the p53 down-stream target genes, *Zmat3*, an inhibitor of cell growth^83^, and *Phlda3*, a repressor of Akt signaling^84^. Commonly down-regulated genes in hyperoxia included microRNA (miR) miR6236, *Sox7*, a TF that promotes EC fate and arterial specification^85^, *Pfkfb3*, a gene important in fructose metabolism that promotes angiogenesis^86^, and *Klhl2*, a regulator of WNK kinases^87^.

However, in addition to this common response, hyperoxia also induced distinct transcriptomic alterations in select EC subtypes, including the CAP1 and CAP2 and proliferative EC (Figure 6F). In early CAP1, hyperoxia suppressed numerous genes that regulate EC homeostasis and angiogenesis (Figure 6F and Table S2). One of the most down-regulated genes was *Ntrk2*, encoding a neurotrophin receptor that promotes neovascularization^88^, EC migration^89^, and vascular relaxation^90^. Hyperoxia also broadly suppressed additional angiogenic genes including *Sparcl1, Glul*^91^, *Ets1*, and *Wfdc1*^92^, and down-regulated MCH class I genes including *H2-Q7*, *H2-K1*, and *H2-D1*. Up-regulated genes included *Lyve1*, a hyaluronan receptor highly expressed by lymphatic EC that promotes dendritic cell trafficking^93^, *Serpine2*, a potent thrombin inhibitor^94^, and the vasoconstrictor *Edn1*. Of note, hyperoxia also increased the expression of *Apln* and *Mmrn2* in the early CAP1, two genes primarily expressed by CAP2 EC under control conditions.

Hyperoxia dysregulated similar pathways, but distinct genes, in proliferative EC (Figure 6F and Table S3). Similar to the effects in early CAP1, hyperoxia suppressed the expression of MHC class I genes. Further, both *Kit* and *Aplnr*, receptors for pathways that promote EC proliferation were specifically suppressed by hyperoxia. Up-regulated genes included *Bax* a gene promoting apoptosis, *Btg2*, a gene that inhibits proliferation, *Gdf15*, a cytokine up-regulated with oxidative stress, *Gtse1*, a gene involved in p53-mediated cell cycle arrest, and *Terf1*, a telomere binding protein that negatively regulates telomere length. Taken together, these data suggest a number of mechanisms that likely contribute to the impaired vascular growth observed in response to hyperoxia.

Hyperoxia also induced unique alterations to the CAP2 transcriptome (Figure 6F and Table S4). Top genes suppressed by hyperoxia included *Cav2*, encoding a protein that prevents abnormal EC proliferation^95^, *Mapt*, a regulator of EC barrier function^96^, and *Hpgd*, important for prostaglandin PGE2 metabolism. Up-regulated genes included *Cd63*, a transmembrane glycoprotein that enhances VEGFR2 downstream signaling^97^, potentially representing a compensatory response given the dependence of CAP2 on epithelial-derived VEGF for survival^31^. Additional up-regulated genes included genes that inhibit NFκB activation (e.g. *Nrros*^98^ and *Tspan6*^99^), a pathway essential for postnatal pulmonary angiogenesis^72^, and a marked increase in *Inhba*, encoding a TGF-β family member that impairs VEGF-mediated EC angiogenesis^100^.

We validated the dysregulated expression of select genes by FISH. In normoxia-exposed P7 mice, numerous MEC in the distal lung exhibited high expression of *Ntkr2* (Figure 6G), in contrast to virtually absent endothelial *Ntrk2* expression in the hyperoxia-exposed lung. In contrast, in normoxia-exposed mice, *Nrros* expression in CAP2 was minimal, but significantly increased by hyperoxia (Figure 6H). A similar pattern was observed with the antiangiogenic gene *Inhba*, with minimal expression in normoxia but high expression in CAP2 exposed to hyperoxia (Figure S8), consistent with a recent report ^15^.

Chronic hyperoxia also appeared to dysregulate the coordinated cross-talk apparent in the capillary EC of the developing lung in normoxia. For example, in normoxia, CAP2 specifically express *Apln*, a pathway essential for embryonic angiogenesis^101^ (Figure 6I). In contrast, CAP1, highly express the gene encoding the apelin receptor, *Aplnr* (Figure 6J). However, in response to hyperoxia, a greater percentage of CAP1 expressed *Apln* (16% vs. 63%), and mean expression increased 4.6-fold (56 vs. 258 cpm). We confirmed this result in lung sections taken from separate cohorts of P7 mice exposed to either normoxia or hyperoxia. Under control conditions, *Apln* was restricted to the CAP2 EC, with multiple *Apln+Car4+* CAP2 present throughout the distal lung. In contrast, minimal *Apln* expression was observed in the CAP1 EC marked by *Plvap* (Figure 6K). Hyperoxia dysregulated this coordinated *Apln* expression, inducing a marked increase in *Apln* expression in multiple *Plvap+* CAP1 EC (arrows). Taken together, these results indicate that hyperoxia had pleiotropic effects on EC development, altering EC subtype abundance and communication, and inducing both shared and subtype-specific changes in gene expression.

### Chronic hyperoxia selectively induces proliferation of venous EC

The presence of a new cluster of proliferating venous EC was intriguing given that the induction of venous EC proliferation in hyperoxia has not been previously described. Moreover, hyperoxia mildly suppressed proliferation in non-venous EC, and markedly decreased both pericyte and myofibroblast proliferation^102^, suggesting that hyperoxia activates unique signaling pathways in venous EC. We first confirmed this computational finding by quantifying proliferating venous EC *in situ* by detecting the venous EC marker, *Slc6a2*, in combination with the proliferation gene *Mki67* in separate cohorts of normoxia- and hyperoxia-exposed P7 pups (Figure 7A). Although we observed proliferating cells located around veins in normoxia-exposed pups, double positive *Slc6a2+Mki67+* proliferating venous EC were extremely rare. In contrast, hyperoxia induced a 7-fold increase in proliferating venous EC (4.4±1.5 vs. 31.3±3.5%, p<0.0001), evidenced by the presence of multiple EC expressing both *Slc6a2* and *Mki67* (arrows) in pulmonary veins (Figure 7B).

**Figure 7:**
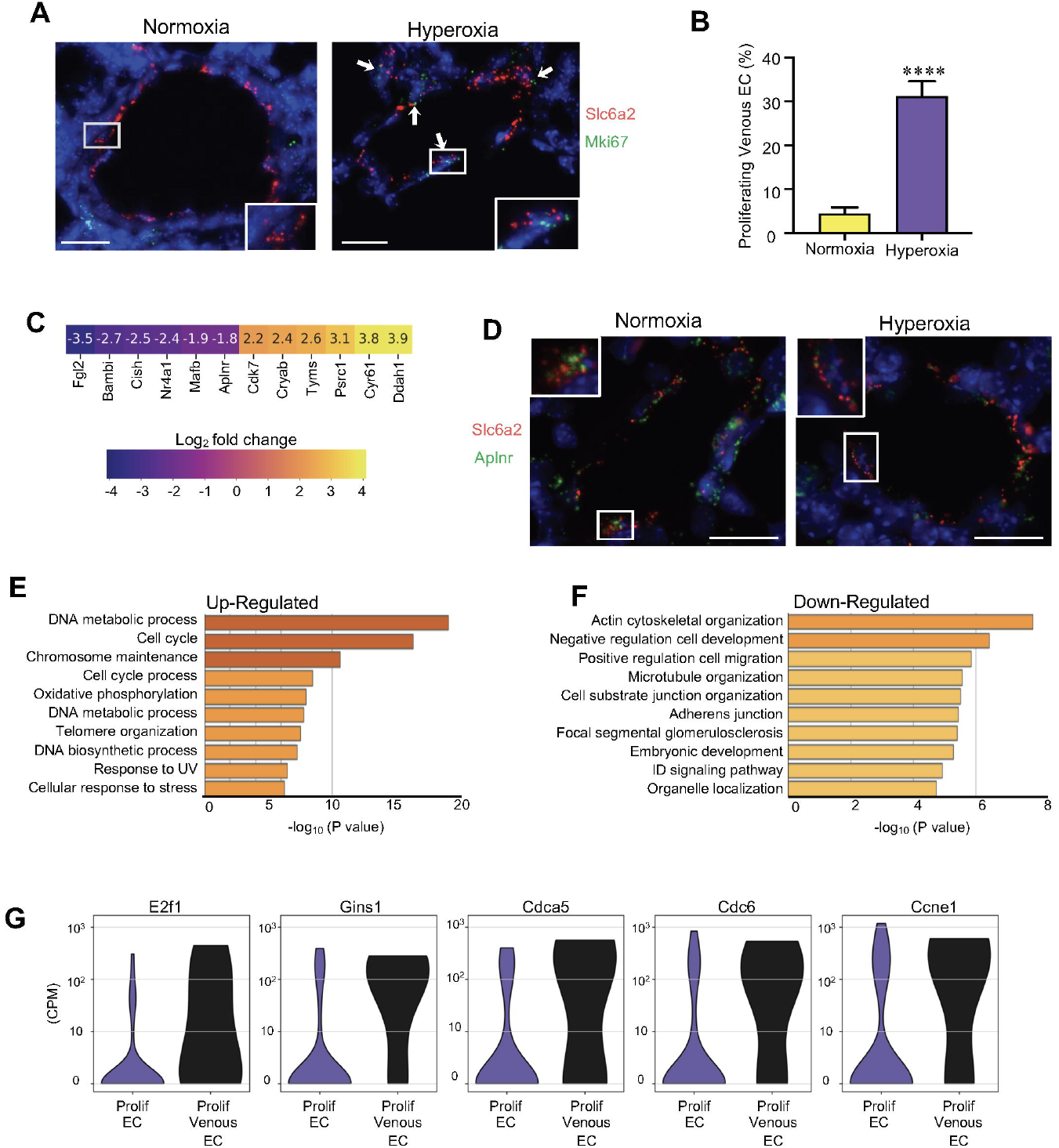
Chronic hyperoxia selectively induces proliferation of venous EC. (A) Representative multiplex *in situ* hybridization to detect *Slc6a2* (red) and *Mki67* (green) in combination with DAPI (blue) in lung sections from P7 mice exposed to either normoxia or hyperoxia. Calibration bar=20μm. (B) Quantification of proliferating venous EC, with data presented are mean±SEM with n=18 veins (normoxia) and n=17 veins (hyperoxia) counted from 4 separate mice in each group, and ****P<0.0001 vs. normoxia via unpaired Student’s t-test. (C) Heat map of select up- and down-regulated genes in hyperoxia versus normoxia at P7 in the venous EC. Values in each square represent the log2 fold changes in average expression. (D) Representative multiplex *in situ* hybridization to detect *Slc6a2* (red) and *Aplnr* (green) in combination with DAPI (blue) in lung sections from P7 mice exposed to either normoxia or hyperoxia. Calibration bar=20μm. (E-F) Pathway analysis of DEG in the proliferating venous EC versus proliferating non-venous EC after excluding venous-specific genes. (G) Violin plots depicting the gene expression of genes highly expressed by proliferating venous EC compared to proliferating non-venous EC.

To further define the hyperoxia-induced alterations in the venous EC transcriptome, we identified differentially expressed genes in venous EC exposed to hyperoxia (Figure 7C). Up-regulated genes included *Ddah1*, which promotes nitric oxide-mediated EC proliferation^103^ and cell cycle regulators such as *Cyr61* and *Cdk7*. Down-regulated genes included the transcription factor *Mafb*, previously implicated in lymphatic patterning^104^, *Bambi*, an inhibitor of TGFβ signaling, and the apelin receptor, *Aplnr*. We validated the decreased *Aplnr* expression by FISH to detect *Slc6a2* in combination with *Aplnr*, and found a high expression of *Aplnr* in venous EC under basal conditions (Figure 7D, left panel), which was decreased in venous EC exposed to hyperoxia (Figure 7D, right panel).

The induction of venous EC proliferation in hyperoxia despite a reduction in *Aplnr*, a key pathway regulating EC proliferation in other models^105^, suggested that alternative pathways were driving venous EC proliferation in response to hyperoxia. To begin to define these distinct mechanisms, we compared the DEGs in proliferating non-venous EC versus proliferating venous EC, after excluding genes that were different between venous EC and all other EC subtypes. We then performed pathway analysis on these remaining DEGs, and found that down-regulated genes in the proliferating venous EC were enriched in pathways related to cytoskeletal rearrangement, migration, and cellular adhesion. In contrast, up-regulated genes were enriched in pathways related to cell cycle regulation, chromosome and telomere maintenance, DNA repair, and cellular response to stress (Figure 7F). Cell cycle regulators more highly expressed in proliferating venous versus non-venous EC included *E2f1, Gins1, Cdca5, Cdc6, and Ccne1* (Figure 7G). Taken together, these data highlight the diverse response of pulmonary EC subtypes to injury, and specifically identify the unique transcriptomic and phenotypic response of venous EC to hyperoxia.

## Discussion

As the lungs transition from a liquid-filled, relatively hypoxic environment *in utero* to air-breathing life, the pulmonary vasculature undergoes dramatic changes to increase blood flow to the lungs and supply oxygen to the body. Subsequently, exponential growth of the microvasculature during early postnatal life drives alveolarization to markedly increase gas exchange surface area. Here, we employed single cell transcriptomics, mechanistic studies in primary EC, and *in situ* hybridization to define the transcriptomic ontogeny of the pulmonary endothelium and the effects of injury on pulmonary EC phenotype during this critical period of vascular growth and remodeling. Our studies highlight the marked increase in EC heterogeneity after birth, the distinction of perinatal lung EC from their adult counterparts, and continued dynamism of the endothelium transcriptome over the first three weeks of life. We show that hyperoxia profoundly altered the perinatal pulmonary endothelium, induced common and unique gene signatures among EC subtypes, stimulated a specific increase in venous EC proliferation, and dysregulated cross-talk between CAP1 and CAP2 EC. Taken together, these data highlight the continued evolution of lung EC phenotype across early postnatal life and the differential sensitivity of the perinatal pulmonary endothelium to injury.

One important finding of our study was the identification of distinct phenotypes of perinatal lung EC compared to adult lung EC, and the presence of a transient arterial subtype not previously identified in recent single cell atlases ^14, 15, 106^. These Art1EC peaked in abundance prior to birth and disappeared by three weeks of life, a time when angiogenesis is slowing and the vasculature is moving toward quiescence. In contrast to Art2 EC which were transcriptionally more similar to adult arterial EC and localized to larger arteries, Art1 EC localized to the distal lung adjacent to capillary EC. Consistent with this finding, Art1 EC embedded between, and shared regulons with both Art2 EC and CAP1, suggesting an intermediate phenotype. This subtype was enriched in numerous genes related to vascular growth and patterning suggesting an important developmental function. Our data demonstrating that by P7, the arterial EC located in the distal lung parenchyma were now expressing Art2 markers suggest that the Art1 EC mature into Art2 EC during the first week of postnatal life. Future mechanistic studies will be important to determine if this temporary arterial subtype plays a role in the rapid microvascular growth coinciding with bulk alveolarization.

Among the microvascular EC, CAP1 spanned three transcriptionally distinct clusters, each peaking at a distinct timepoint. This dynamic diversity of phenotype across development challenges recent descriptions of CAP1 as a static, general capillary that resembles capillaries in other organs^12^. Genes differentially expressed by CAP1 during early alveolarization suggest that they exhibit greater proliferative and migratory potential than their late counterparts. This is consistent with our prior work demonstrating that primary microvascular EC isolated from the early alveolar lung proliferate and migrate more quickly than those isolated from the adult^72^. Prior reports have suggested that a subset of CAP1 MEC may be capable of self-renewal^106^. Consistent with this notion, we identified *Peg3* as highly expressed by early CAP1, a finding not previously highlighted in prior reports. *Peg3* is expressed in specific progenitor populations capable of self-renewal in adult tissues^68^, and *Peg3*+ EC are abundant in the embryonic vasculature and appear to mark EC with high proliferative potential^107^. By silencing *Peg3* in primary EC isolated from the early alveolar lung we demonstrated that *Peg3* is required for CAP1 proliferation. However, definitive studies determining whether *Peg3* confers a capacity for the CAP1 to self-renew need to be performed. Of note, most proliferative EC lacked *Car4*, and many of them expressed *Peg3*, supporting the notion that proliferating EC form a subset of CAP1. Future studies to selectively delete *Peg3* from lung EC will be important to determine if *Peg3* positive CAP1 serve to replenish the capillary EC niche, as has been observed in other tissues. *Peg3* also promotes nuclear factor kappa-B (NFκB) activation^108^, a pathway we have shown is essential for angiogenesis during early alveolarization^70, 72^. Our studies in human lung tissue demonstrated, for the first time, that *PEG3* expression is high in human lung microvascular EC during early alveolarization, but diminishes as alveolarization ends. Taken together, these data suggest that developmental expression of *Peg3* in CAP1 may represent an evolutionarily conserved pathway to drive the expansion of the pulmonary microcirculation during this early phase of postnatal distal lung growth.

Our data also revealed dynamic progression of the CAP2 phenotype with maturation, and identified putative novel functions for this subtype unique to the developing lung. Although a recent report identified a very small number of CAP2 present in the lung at E18^106^, abundance markedly increased by P0, consistent with our findings. Further, the expression of multiple genes that distinguish CAP2 (e.g. *Car4, Fibin*) increase during early postnatal life. Taken together, these data suggest that although CAP2 speciation may begin just prior to birth, the physiologic changes occurring after birth (e.g. increased oxygen tension, shear stress, cyclic stretch) may represent important environmental signals driving this distinct phenotype. During late sacculation (P1), aerocytes highly expressed genes related to ameboid cell migration and cytoskeletal rearrangement, perhaps reflecting dynamic extension of the multiple projections extending from these EC to encircle the alveolus^12^. By early alveolarization, genes related to vascular development increased, suggesting a putative novel function for CAP2 in promoting the angiogenesis that drives alveolarization. By late alveolarization, genes important for antigen presentation and response to interferon were up-regulated. The heightened expression of immune response genes in combination with the location of these cells abutting the alveolus suggest that CAP2 may subsume a greater role in immunity and response to inhaled pathogens as the lung matures.

Ongoing growth of the lung after birth increases the susceptibility to injuries that disrupt lung development. In our study, hyperoxia induced multiple alterations in the endothelial compartment. First, hyperoxia induced a new cluster of proliferating venous EC. This finding is potentially consistent with a recent report showing that although hyperoxia did not alter EC proliferation as a whole, it did increase the number of vWF+ proliferating EC ^109^. Given that *Vwf* is most highly expressed by venous EC (and not expressed by capillary EC), it is likely that a component of the proliferating EC in that study were venous EC. This finding is intriguing, given that venous EC in the brain and retina were recently shown to proliferate, migrate against blood flow, and differentiate into tip, capillary and arterial EC ^110^. Moreover, a subset of infants born prematurely who develop the disrupted angiogenesis characteristic of BPD, develop pulmonary vein stenosis, a complication associated with high mortality, and molecular mechanisms that remain poorly defined^111^. However, whether the induction of venous EC proliferation by hyperoxia serves a compensatory versus pathologic role is not clear, and will be an important question for future studies. Second, hyperoxia induced a common gene signature across all EC characterized broadly by an up-regulation of genes that suppress cell growth and proliferation, and down-regulation of genes that promote angiogenesis and EC fate, providing novel potential mechanisms for the impaired angiogenesis observed in the hyperoxia-exposed lung. Third, hyperoxia induced distinct alterations in select microvascular EC subtypes, including a suppression of MHC class I expression in both CAP1 and proliferating MEC, suggesting an impaired adaptive immune responses or induction of local tolerance in the hyperoxic lung^112^. This finding may have important clinical implications, as lung infections represent a source of secondary injury in patients with BPD, further increasing respiratory morbidity. Fourth, the induction of *Apln* expression in hyperoxia-exposed CAP1 is interesting given that this gene is normally restricted to CAP2. This may represent a significant limitation of the use of *Apln*-CreERT2 mice to lineage trace CAP2 during injury^12^. Future studies to lineage trace CAP1 to determine if *Apln* induction is an early sign of differentiation into CAP2 after injury will be important next steps. Finally, hyperoxia slightly increased the abundance of early CAP1 while decreasing the abundance of late CAP1, consistent with the arrest in lung development apparent histologically.

Our study builds upon recent reports characterizing lung cell heterogeneity across development, and the response of coarse cell types (epithelial, endothelial, mesenchymal, and immune) to injury. However, by focusing on a single cell type, we were able to delineate the transcriptome of the immature endothelium, the developmental evolution of endothelial phenotype, and the pleiotropic effects of hyperoxia at a greater resolution than previously described. Additionally, in our preliminary studies, we found that lung EC survival was markedly impaired by commonly employed semi-automated mechanical dissociators^7, 15^, potentially causing loss of more vulnerable EC subtypes in prior publications that utilized this method of dissociation. Finally, the higher sequencing depth we achieved with plate-based rather than droplet-based technology may have permitted a greater capacity to detect differences in the expression of low abundance transcripts^113^.

However, our study also has a number of limitations. First, marked changes occurred in the transcriptome between our final two timepoints, precluding our ability to identify discrete transcriptomic and phenotypic alterations occurring within this developmental window. Second, there are some limitations to computational identifications of TF activity from RNA sequencing data. We were not able to infer post-transcriptional regulatory interactions, TFs with unknown binding motifs, or compare regulatory networks against each other, only across cells^114^. Third, our data only included a single hyperoxic timepoint, limiting our capacity to elucidate early alterations that may underlie the phenotype observed by P7, or durable alterations in cell phenotype after recovery. Finally, the inclusion of a single male and female mouse per timepoint limited our ability to identify potential sex-dependent transcriptomic differences during development and injury. This represents an important area for future research given the well-known sexual dimorphism observed in both the incidence and severity of BPD, as well is in the vascular response to hyperoxic injury observed in preclinical studies^115^.

In summary, this study is the first to provide an in-depth delineation of pulmonary EC during a period of rapid postnatal lung growth. Our data highlight the unique phenotypes of perinatal pulmonary EC at both the macro and microvascular level, and comprehensively identify potential genes, pathways, and EC subtypes driving postnatal development of the pulmonary circulation and the progression towards quiescence. Further, our data provides a substantially more granular report of the response of the pulmonary endothelium to hyperoxic injury than previously published, including the identification of select EC subtypes, especially venous EC, with a heightened susceptibility and unique response to injury. Taken together, our results and interpretation provide an essential framework for understanding how the immature pulmonary endothelium differs from the mature endothelium at single cell resolution, with important implications for diseases characterized by impaired lung and vascular growth, pathologic vascular remodeling, and lung injury.

## Acknowledgements

We thank Sai Saroja Kolluru (Stanford University) for assistance with library submission to the Chan Zuckerberg Biohub and to David Sassoon and Giovanna Marazzi (UCSF) for the providing us with the anti-Peg3 antibody. This work was supported by National Institutes of Health grants HL122918 (CMA), HL 154002 (CMA), HL155828 (CMA), HD092316 (CMA, DNC), HL160018 (CMA, DNC) the Stanford Maternal Child Health Institute Tashia and John Morgridge Faculty Scholar Award (CMA), the Crandall Endowed Faculty Scholar Award (CMA), the Stanford Center of Excellence in Pulmonary Biology (DNC), Bill and Melinda Gates Foundation (SRQ), the Chan Zuckerberg Biohub (DNC and SRQ), the Ernest and Amelia Gallo Endowed Fellowship (NES), and the Chan Zuckerberg Biohub Physician-Scientist Fellowship (NES).

## Author contributions

Conceptualization, C.M.A., D.N.C. and S.R.Q.; Methodology, F.Z. and S.R.Q.; Software, F.Z. and C.K.; Formal Analysis, F.Z., S.H.D., C.K., and C.M.A.; Investigation, X.C., M.L., R.D.G., N.E.S., D.Z., and R.C.J.; Resources, G.H.P.; Data Curation, F.Z. and C.K., Writing-Original Draft, C.M.A. and F.Z.; Writing-Review & Editing, C.M.A., F.Z. and D.N.C. Supervision, C.M.A.; Project Administration, C.M.A. and D.N.C.; Funding Acquisition, C.M.A., D.N.C. and S.R.Q.

## Methods

### Mouse lung cell isolation

C57BL/6 mice were obtained from Charles River Laboratories. For studies using E18.5, P1, and P7 murine lungs, pregnant dams were purchased, and pups aged prior to lung isolation. At E18.5, dam was asphyxiated with CO2 and pups extracted. At P1, P7, and P21 pups were euthanized with euthanasia solution (Vedco Inc.). Genetic sex of mice at developmental stages E18.5 and P1 was determined by performing PCR amplification of the Y chromosome gene Sry. P7 and P21 mice were sexed through identification of a pigment spot on the scrotum of male mice^116^. For all timepoints, single female and male mice were randomly selected for the studies. For all timepoints, except E18.5, the pulmonary circulation was perfused with ice cold heparin in 1x PBS until the circulation was cleared of blood. Lungs were minced and digested with Liberase (Sigma Aldrich) in RPMI for 15 (E18.5, P1, and P7) or 30 (P21) minutes at 37C, 200 rpm. Lungs were manually triturated and 5% fetal bovine serum (FBS) in 1x PBS was used to quench liberase solution. Red blood cells were lysed with 1x RBC lysis buffer (ThermoFisher) as indicated by the manufacturer and total lung cells counted on BioRad cell counter (BioRad). Protocols for the murine studies adhered to American Physiological Society/US National Institutes of Health guidelines for humane use of animals for research and were prospectively approved by the Institutional Animal Care and Use Committee at Stanford (APLAC #19087).

### Chronic Hyperoxia

Newborn C57BL/6 mice pups at P0 were maintained in 80% O2 (hyperoxia) in a BioSpherix chamber (BioSpherix, Parish, NY) for 7 days, an established experimental model that recapitulates the impaired alveolarization observed in BPD^117^. Dams were rotated every 24h to prevent oxygen toxicity. Pups were euthanized at P7 and single lung cells obtained as described above.

### Latex Dye Injection of the Pulmonary Circulation

Mice were anesthetized with pentobarbital and the chest cavity opened. While the heart was still beating, blue microfil dye (Cat, LMV120, Flow Tek Inc.) was injected into right ventricle slowly and steadily using a 26-gauge 1-ml syringe needle. After allowing the dye to solidify, the lungs were removed, washed in PBS briefly and fixed in 4% PFA overnight. For whole mount imaging, the lungs were then dehydrated in methanol twice and cleared with 1:1 benzyl alcohol/benzyl benzoate (Sigma-Aldrich), and photos of the lungs taken next to a 10 cm ruler to allow inclusion of calibration bar on the images.

### Immunostaining and fluorescence-activated cell sorting (FACS) of single cells

Lung cells were plated at 1×10^6^ cells per well and stained with Fc block (CD16/32, 1:100, Tonbo Biosciences) for 30 min on ice. Cells were surface stained with the endothelial marker CD31 (1:100, clone: MEC3.1, eBiosciences), epithelial marker CD326 (1:100, clone: CD326, eBiosciences), and immune marker CD45 (1:100, clone: F11, eBiosciences) for 30 min on ice. The live/dead dye, SYTOX Blue (ThermoFisher), was added to cells and incubated for 3 min prior to sorting into 384-well plates (Bio-Rad Laboratories, Inc) prefilled with lysis buffer using the Sony LE-SH800 cell sorter (Sony Biotechnology Inc), a 100μm sorting chip (Catalog number: LE-C3110) and ultra-purity mode. Single color controls were used to perform fluorescence compensation and generate sorting gates. 384-well plates containing single cells were spun down, immediately placed on dry ice and stored at −80C.

### cDNA library generation using Smart-Seq2

Complementary DNA from sorted cells was reverse transcribed and amplified using the Smart-Seq2 protocol on 384-well plates as previously described^118, 119^. Concentration of cDNA was quantified using picogreen (Life technology corp.) to ensure adequate cDNA amplification. In preparation for library generation, cDNA was normalized to 0.4 ng/uL. Tagmentation and barcoding of cDNA was prepared using in-house Tn5 transposase and custom, double barcoded indices^6^. Library fragment concentration and purity were quantified by Agilent bioanalyzer. Libraries were pooled and sequenced on Illumina NovaSeq 6000 with 2×100 base kits and at a depth of approximately 1 million paired reads per cell.

### Data analysis and availability

Sequencing reads were mapped against the mouse genome (GRCm38) using <u>STAR aligner</u> and genes were counted using <u>HTSeq</u>^120, 121^. To coordinate mapping and counting on Stanford’s high-performance computing cluster, <u>snakemake</u> was used^122^. Gene expression count tables were converted into loom objects (https://linnarssonlab.org/loompy/) and cells with less than 50,000 uniquely mapped reads or less than 400 genes per cell were discarded. Doublets were removed in three ways. First, clusters that expressed epithelial, endothelial or mesenchymal genes were excluded. Clusters showing joint expression of mutually exclusive cell type markers (e.g. *Epcam, Ptprc, Col6a2*) with *Cdh5* were manually excluded as well. Finally, a few remaining doublets were removed manually. Counts for the remaining 2931 cells were normalized to counts per million reads. For Uniform Manifold Approximation and Projection (UMAP)^16^, 500 features were selected that had a high Fano factor in most mice, and the restricted count matrix was log-transformed with a pseudocount of 0.1 and projected onto the top 25 principal components using <u>scikit-learn</u>^123^. Unsupervised clustering was performed using <u>Leiden</u>^17^. Batch-corrected KNN^124^ was used to compare our data with Tabula Muris^6^. Singlet (https://github.com/iosonofabio/singlet) and for specific analyses, custom Python 3 scripts were used: the latter available at https://github.com/iosonofabio/lung_neonatal_endothelial. Pathway analysis was performed using Metascape^125^. Pseudotime was determined with Scanpy: https://scanpy.readthedocs.io/en/stable/api/scanpy.tl.dpt.html?highlight=pseudotime#scanpy.tl.dpt. RNA velocity was performed using scVelo^57^, which explicitly models curly developmental trajectories^126^. SCENIC was used to perform gene regulatory network inference using the combined datasets of GSE147668, GSE159804, and GSE172251^32^. Data from Tabula Muris^78^ was downloaded from Figshare. Raw fastq files, count tables, and metadata are available on NCBI’s Gene Expression Omnibus (GEO) under submission GSE159804. https://www.ncbi.nlm.nih.gov/geo/query/acc.cgi?acc=GSE159804.

#### Human lung tissue

De-identified, formalin-fixed, paraffin-embedded postmortem lung tissue was provided by the NHLBI LungMAP Human Tissue Core Biorepository (BRINDL). The Biorepository is approved by the University of Rochester Research Subjects Review Board (RSRB00056775). Informed consent for research use has been provided for each sample. Clinical metadata and histopathology are listed in Table S5.

### In-situ validation using RNAscope and immunofluorescence (IF)

Embryonic and post-natal mice were euthanized as described above. Female and male mice were randomly selected from the litter, and at least 2 litters were used to source the lung tissue for all validation studies. P1, P7, and P21 murine lungs were perfused as described above, and P7 and P21 lungs inflated with 2% low melting agarose (LMT) in 1xPBS and placed in 10% neutral buffered formalin. Following 20 hours incubation at 4C, fixed lungs were washed twice in 1xPBS and placed in 70% ethanol for paraffin-embedding. In situ validation of genes identified by scRNAseq was performed using the RNAscope Multiplex Fluorescent v2 Assay kit (Advanced Cell Diagnostics) and according to the manufacturer’s protocol. Formalin-fixed paraffin-embedded (FFPE) lung sections (5 μm) were used within a day of sectioning for optimal results. Nuclei were counterstained with DAPI (Life Technology Corp.) and extracellular matrix proteins stained with hydrazide^127^. Opal dyes (Akoya Biosciences) were used for signal amplification as directed by the manufacturer. Images were captured with Zeiss LSM 780 and Zeiss LSM 880 confocal microscopes, using 405nm, 488nm, 560nm and 633nm excitation lasers. For scanning tissue, each image frame was set as 1024×1024 and pinhole 1AiryUnit (AU). For providing Z-stack confocal images, the Z-stack panel was used to set z-boundary and optimal intervals, and images with maximum intensity were processed by merging Z-stacks images. For all both merged signal and split channels were collected.

### Quantification of RNA Scope Images

For select RNA scope experiments, the number and location of specific cells were quantified. Adjustments to the brightness and contrast of an image to increase the clarity of the signal was always applied to the entire image and never to separate areas of the image. For the determination of the percentage of *Peg3*^+^ cells located in the distal lung parenchyma, at least 4 separate images were quantified per mouse. For the determination of proliferating venous EC, 4-5 veins per mouse were imaged and quantified. For all quantification analysis, a cell was deemed positive if it contained >1 puncta per cell for a specific gene.

### Isolation of Primary Lung Microvascular EC

PEC were isolated from P7 C57BL/6 mice as described previously^72^. Briefly, primary MEC were obtained from 10-12 mouse lungs by digestion of peripheral lung tissue with collagenase IA (0.5 mg/ml, Sigma, St. Louis, MO) for 30 min at 37°C, followed by incubating cell lysate for 15 min at RT with anti-CD31-coated magnetic beads (Dynabeads, Invitrogen, Carlsbad, CA), resulting in between 2-3 million MEC after bead selection. MEC isolated by this method were previously characterized by flow cytometry and found to be exclusively CD45-, greater than 95% positive for the endothelial specific marker, CD102 (E2), and greater than 80% demonstrating binding to Griffonia simplicifolia, indicating primarily microvascular EC. Primary MEC were cultured in endothelial growth media (EGM) containing 5% FBS with growth factors (EBM-2; Lonza, Basel, Switzerland) at 37°C in 5% CO_2_. PEC from passage 0-2 were used for all experiments.

### RNA Interference

Primary lung MEC isolated from P7 mice were transfected with 25 nM of NTC, or Peg3 On-Target Plus SMART pool siRNA (Dharmacon, Lafayette, CO; Thermo Fisher Scientific, Lafayette, CO) using Lipofectamine 2000 (Invitrogen) for 6h as previously described^69^. The groups of MEC were then recovered for 42h, prior to use for proliferation assays and immunocytochemistry.

### Proliferation Assays

Proliferation of siRNA transfected primary lung MEC proliferation was determined by BrdU incorporation assays. MEC (6 × 10^3^) were plated into each well of a 96-well plate and synchronized with starvation media (0.2% FBS) for 16h, prior to stimulation with experimental media. The incorporation of BrdU was then measured by ELISA at 24h per the manufacturer’s protocol (Roche Diagnostics, Mannheim, Germany).

### Statistical Analyses

To identify differentially expressed genes within cell populations, Kolmogorov Smirnov tests on the distributions of gene expression were performed on all genes, and the genes with the largest test statistic (lowest P value) were chosen as the top differentially expressed genes. Data are presented as mean ± SD. Differences between two groups was determined by Student’s t-test and either one-way or two-way ANOVA for detecting differences between more than 2 groups. For all analyses, a P value of ≤0.05 considered statistically significant.

