## Supplemental Table 5 for "Developmental diversity and unique sensitivity to injury of lung endothelial subtypes during a period of rapid postnatal growth"

| **Table S5: Clinical Metadata from Human Lung Tissue** | | | |
| --- | --- | --- | --- |
| **Donor** | **Age at Death** | **Cause of Death** | **Histopathology** |
| Donor 1 | 1 Day | Anencephaly | Normal lung growth and alveolar structure |
| Donor 2 | 6 months | Blunt Injury/MVA | Normal lung development and structure |
| Donor 3 | 3 years | Brain injury | Normal lung growth and structure. Rare mucus aspiration. Focal acute adventitial hemorrhage. |
| Donor 4 | 24 years | Probable poisoning | Normal growth. Rare focus of bronchiolar remodeling. |
