## Supplemental Table 4 for "Developmental diversity and unique sensitivity to injury of lung endothelial subtypes during a period of rapid postnatal growth"

| **Table S4: Top Up- and Down-regulated Genes in the aCAP in Hyperoxia vs. Normoxia** | | | |
| --- | --- | --- | --- |
| **Up-Regulated Genes in Hyperoxia** | | | |
| **Gene** | **P-value** | **Log2 Fold Change** | **Statistic** |
| Rtl6 | 3.53E-11 | 5.96 | 0.61 |
| Cd63-ps | 0.0043 | 4.78 | 0.31 |
| Cd63 | 0.0043 | 4.77 | 0.31 |
| Snhg15 | 0.00180 | 4.62 | 0.33 |
| Eda2r | 1.56E-05 | 4.41 | 0.42 |
| Uap1l1 | 0.00011 | 4.02 | 0.39 |
| Inhba | 1.01E-06 | 3.97 | 0.47 |
| Mir99ahg | 0.00086 | 3.28 | 0.34 |
| Gpx3 | 9.37E-06 | 3.13 | 0.43 |
| Ahnak | 0.00020 | 3.03 | 0.38 |
| Zfp862-ps | 0.00360 | 2.78 | 0.31 |
| Zmat3 | 0.00016 | 2.73 | 0.38 |
| Tm4sf1 | 0.00228 | 2.52 | 0.32 |
| Sdc4 | 7.99E-05 | 2.47 | 0.39 |
| Spry2 | 0.00040 | 2.47 | 0.36 |
| Olfr613 | 0.00377 | 2.45 | 0.31 |
| Itpripl1 | 3.99E-06 | 2.43 | 0.45 |
| Nrros | 8.24E-06 | 2.41 | 0.44 |
| Zfa-ps | 1.89E-05 | 2.37 | 0.42 |
| Ccdc36 | 0.00217 | 2.35 | 0.32 |
| Tspan6 | 2.02E-06 | 2.27 | 0.46 |
| Gap43 | 1.38E-05 | 2.20 | 0.43 |
| Plekhh2 | 1.62E-08 | 2.18 | 0.53 |
| Cdkn1a | 2.96E-13 | 2.13 | 0.66 |
| Aldoart1 | 0.003604 | 2.08 | 0.31 |
| **Down-Regulated Genes in Hyperoxia** | | | |
| **Gene** | **P-value** | **Log2 Fold Change** | **Statistic** |
| Dennd2d | 0.00074 | -3.49 | 0.35 |
| D630045J12Rik | 0.00028 | -2.41 | 0.37 |
| Mir6236 | 1.21E-12 | -2.34 | 0.64 |
| Slc16a9 | 0.00027 | -2.32 | 0.37 |
| Cav2 | 8.51E-05 | -2.28 | 0.39 |
| Txnip | 0.00026 | -2.27 | 0.37 |
| Lrp10 | 0.00218 | -2.23 | 0.32 |
| Pllp | 3.29E-05 | -2.20 | 0.41 |
| Sox7 | 0.00368 | -2.13 | 0.31 |
| Lars2 | 2.49E-08 | -2.11 | 0.52 |
| Dpp4 | 0.00016 | -1.77 | 0.38 |
| Cyth3 | 0.00117 | -1.56 | 0.34 |
| Mapt | 0.00116 | -1.45 | 0.34 |
| Rtn1 | 6.29E-05 | -1.43 | 0.40 |
| Fgf16 | 2.32E-09 | -1.43 | 0.56 |
| Hpgd | 2.63E-08 | -1.40 | 0.52 |
| Pard6g | 0.00381 | -1.38 | 0.31 |
| Tbxa2r | 0.002331 | -1.37 | 0.32 |
| Mest | 1.89E-06 | -1.37 | 0.46 |
| 1700066B19Rik | 1.96E-08 | -1.36 | 0.53 |
| Rsrp1 | 0.00117 | -1.30 | 0.34 |
| Nectin3 | 0.00023 | -1.27 | 0.37 |
| Slco2a1 | 0.00202 | -1.20 | 0.33 |
| Gatad2a | 0.00406 | -1.18 | 0.31 |
| Bcam | 0.00289 | -1.17 | 0.32 |
