## Supplemental Table 3 for "Developmental diversity and unique sensitivity to injury of lung endothelial subtypes during a period of rapid postnatal growth"

| **Table S3: Top Up- and Down-regulated Genes in the Proliferative EC in Hyperoxia vs. Normoxia** | | | |
| --- | --- | --- | --- |
| **Up-Regulated Genes in Hyperoxia** | | | |
| **Gene** | **P-value** | **Log2 Fold Change** | **Statistic** |
| Gdf15 | 0.00027 | 7.34 | 0.34 |
| Eda2r | 4.93E-06 | 6.88 | 0.40 |
| Dcxr | 4.01E-05 | 6.39 | 0.37 |
| Sulf2 | 0.00015 | 5.97 | 0.35 |
| Dglucy | 0.00087 | 5.91 | 0.31 |
| Zmat3 | 1.64E-07 | 5.64 | 0.45 |
| Rtl6 | 0.00049 | 4.92 | 0.33 |
| 4930413G21Rik | 0.00051 | 3.35 | 0.32 |
| Ephx1 | 4.03E-07 | 2.88 | 0.44 |
| Galnt17 | 0.00049 | 2.80 | 0.33 |
| Sdc4 | 2.98E-07 | 2.71 | 0.45 |
| n-R5-8s1 | 0.00030 | 2.68 | 0.33 |
| Serpine2 | 5.51E-05 | 2.63 | 0.37 |
| Mospd2 | 0.00130 | 2.62 | 0.31 |
| Tnfrsf10b | 6.82E-05 | 2.54 | 0.36 |
| Gja1 | 0.00034 | 2.50 | 0.33 |
| Rap2b | 7.46E-07 | 2.40 | 0.43 |
| Phlda3 | 3.25E-11 | 2.24 | 0.55 |
| Mmrn2 | 6.34E-11 | 2.19 | 0.55 |
| Bax | 5.12E-10 | 2.09 | 0.52 |
| Bloc1s2-ps | 0.00091 | 1.95 | 0.31 |
| Prnp | 5.28E-06 | 1.95 | 0.40 |
| Cdkn1a | 5.29E-08 | 1.95 | 0.47 |
| Cd44 | 0.00026 | 1.80 | 0.34 |
| Igf2r | 1.48E-05 | 1.68 | 0.39 |
| **Down-Regulated Genes in Hyperoxia** | | | |
| **Gene** | **P-value** | **Log2 Fold Change** | **Statistic** |
| H2-Q6 | 0.00068 | -3.01 | 0.32 |
| H2-Q7 | 1.21E-09 | -2.26 | 0.52 |
| Dennd2d | 0.00084 | -2.26 | 0.31 |
| Glp1r | 0.00078 | -1.91 | 0.32 |
| H2-K1 | 8.88E-10 | -1.82 | 0.52 |
| Ube2m | 0.00020 | -1.77 | 0.34 |
| Mir6236 | 9.99E-16 | -1.75 | 0.68 |
| Slc6a6 | 0.00067 | -1.68 | 0.32 |
| Fhl1 | 0.00012 | -1.67 | 0.35 |
| Glul | 1.85E-05 | -1.63 | 0.38 |
| Aplnr | 0.00055 | -1.62 | 0.32 |
| Tlnrd1 | 0.00137 | -1.51 | 0.30 |
| Csrp1 | 2.48E-06 | -1.47 | 0.41 |
| Aqp1 | 9.25E-05 | -1.37 | 0.36 |
| Arf6 | 5.26E-05 | -1.36 | 0.37 |
| Cav2 | 0.00030 | -1.35 | 0.33 |
| Mdc1 | 6.90E-06 | -1.34 | 0.40 |
| Mtch1 | 1.65E-06 | -1.31 | 0.42 |
| Ppia | 0.00016 | -1.27 | 0.35 |
| Ets1 | 6.44E-06 | -1.27 | 0.40 |
| Kit | 0.00026 | -1.19 | 0.34 |
| Ppp1ccb | 0.00046 | -1.19 | 0.33 |
| Hpgd | 1.01E-06 | -1.12 | 0.43 |
| CAAA01147332.1 | 5.13E-05 | -1.08 | 0.37 |
| Hnrnpb | 2.41E-08 | -1.07 | 0.48 |
