## Supplemental Table 2 for "Developmental diversity and unique sensitivity to injury of lung endothelial subtypes during a period of rapid postnatal growth"

| **Table S2: Top Up-and Down-regulated Genes in the Early gCAP in Hyperoxia vs. Normoxia** | | | |
| --- | --- | --- | --- |
| **Up-Regulated Genes in Hyperoxia** | | | |
| **Gene** | **P-value** | **Log2 Fold Change** | **Statistic** |
| Eda2r | 1.03E-21 | 4.56 | 0.42 |
| Zmat3 | 7.88E-17 | 4.05 | 0.37 |
| Serpine2 | 6.96E-21 | 3.55 | 0.41 |
| Cdkn1a | 4.35E-63 | 3.03 | 0.72 |
| n-R5-8s1 | 1.20E-22 | 3.02 | 0.43 |
| Ephx1 | 4.62E-28 | 2.82 | 0.48 |
| Sdc4 | 4.66E-19 | 2.29 | 0.39 |
| Icam1 | 4.66E-19 | 2.26 | 0.36 |
| Apln | 5.37E-12 | 2.19 | 0.31 |
| Lyve1 | 6.58E-54 | 2.18 | 0.67 |
| Phlda3 | 7.15E-22 | 2.11 | 0.42 |
| Mmrn2 | 3.23E-22 | 2.04 | 0.43 |
| Cyp4b1 | 6.77E-24 | 1.77 | 0.44 |
| Igf2r | 4.83E-21 | 1.70 | 0.42 |
| Bax | 1.38E-34 | 1.65 | 0.53 |
| Prnp | 2.60E-14 | 1.60 | 0.34 |
| Ecm1 | 2.46E-16 | 1.57 | 0.37 |
| Exoc4 | 1.19E-11 | 1.43 | 0.30 |
| Ccnd2 | 6.54E-17 | 1.43 | 0.37 |
| Edn1 | 1.05E-12 | 1.41 | 0.32 |
| Ccnd1 | 2.59E-17 | 1.29 | 0.38 |
| Itpripl1 | 3.29E-12 | 1.27 | 0.31 |
| Myh9 | 2.18E-13 | 1.13 | 0.32 |
| 4933429H19 | 2.77E-12 | 1.12 | 0.31 |
| Slc39a1 | 1.05E-12 | 1.06 | 0.32 |
| **Down-Regulated Genes in Hyperoxia** | | | |
| **Gene** | **P-value** | **Log2 Fold Change** | **Statistic** |
| Ntkr2 | 7.43E-19 | -4.16 | 0.39 |
| H2-Q7 | 4.18E-22 | -2.98 | 0.43 |
| Sparcl1 | 3.52E-16 | -2.13 | 0.36 |
| Pfkfb3 | 1.84E-14 | -1.94 | 0.34 |
| Olfr623 | 4.37E-14 | -1.83 | 0.33 |
| Slc6a6 | 1.31E-20 | -1.75 | 0.41 |
| Wfdc1 | 1.11E-16 | -1.73 | 0.36 |
| Mir6236 | 3.42E-47 | -1.62 | 0.63 |
| Tmeff1 | 1.47E-12 | -1.61 | 0.31 |
| Rn7sk | 1.25E-17 | -1.60 | 0.31 |
| H2-K1 | 4.04E-18 | -1.51 | 0.38 |
| Glul | 2.45E-12 | -1.50 | 0.38 |
| Mest | 2.31E-14 | -1.45 | 0.31 |
| Csrp1 | 1.62E-12 | -1.35 | 0.34 |
| Aqp1 | 4.44E-13 | -1.22 | 0.31 |
| Ets1 | 5.19E-12 | -1.12 | 0.32 |
| Ppia | 2.20E-13 | -1.12 | 0.31 |
| H2-D1 | 3.57E-18 | -0.97 | 0.32 |
| Fgf16 | 4.60E-26 | -0.96 | 0.46 |
| Hnrnpab | 5.01E-12 | -0.95 | 0.31 |
| Gja4 | 6.96E-12 | -0.95 | 0.31 |
| CT010467 | 4.20E-19 | -0.79 | 0.40 |
| Mfap2 | 3.50E-14 | -0.76 | 0.33 |
| Lars2 | 3.72E-17 | -0.74 | 0.37 |
| Ucp2 | 5.32E-12 | -0.74 | 0.31 |
