## Supplemental Table 1 for "Developmental diversity and unique sensitivity to injury of lung endothelial subtypes during a period of rapid postnatal growth"

| **Table S1: Top Differentially-Regulated Genes in the Arterial I vs. Arterial II EC** | | | |
| --- | --- | --- | --- |
| **Up-Regulated Genes in Arterial 1** | | | |
| **Gene** | **P-value** | **Log2 Fold Change** | **Statistic** |
| Col4a2 | 3.14E-07 | 3.25 | 0.62 |
| Gpihbp1 | 5.87E-05 | 3.18 | 0.51 |
| Scn7a | 1.46E-06 | 2.89 | 0.59 |
| Cd36 | 0.00023 | 2.86 | 0.48 |
| Col4a1 | 4.14E-06 | 2.85 | 0.57 |
| Rasgrp3 | 0.00016 | 2.65 | 0.49 |
| Tmem204 | 0.00287 | 2.61 | 0.41 |
| Igfbp7 | 0.00080 | 2.51 | 0.45 |
| Tbx3 | 0.00190 | 2.39 | 0.42 |
| Mest | 0.00027 | 2.39 | 0.48 |
| Ets1 | 0.00050 | 2.28 | 0.46 |
| Emcn | 0.00043 | 2.23 | 0.46 |
| Robo4 | 0.00092 | 2.18 | 0.44 |
| Kit | 0.00058 | 2.05 | 0.45 |
| BC028528 | 1.63E-05 | 2.04 | 0.54 |
| Afap1l1 | 0.00012 | 2.02 | 0.50 |
| Clec1a | 0.00375 | 1.99 | 0.40 |
| C920021L13Rik | 0.00012 | 1.82 | 0.50 |
| Ctla2a | 0.00080 | 1.60 | 0.45 |
| Bmpr2 | 0.00190 | 1.57 | 0.42 |
| Plk2 | 2.86E-05 | 1.55 | 0.53 |
| Aqp1 | 0.00190 | 1.50 | 0.42 |
| Keap1 | 0.00375 | 1.39 | 0.40 |
| Plvap | 9.90E-05 | 1.39 | 0.50 |
| Ivns1abp | 0.00080 | 1.38 | 0.45 |
| **Up-Regulated Genes in Arterial 2** | | | |
| **Gene** | **P-value** | **Log2 Fold Change** | **Statistic** |
| Cytl1 | 0.00050 | 12.87 | 0.46 |
| Lox | 0.00218 | 9.14 | 0.42 |
| Gcnt1 | 0.00050 | 8.65 | 0.46 |
| Rnase1 | 0.00016 | 6.17 | 0.49 |
| Sulf2 | 0.00328 | 5.88 | 0.40 |
| S100a4 | 0.00080 | 5.32 | 0.45 |
| Mgp | 5.55E-16 | 5.00 | 0.89 |
| Trpv4 | 4.11E-05 | 4.49 | 0.52 |
| Cdh13 | 3.94E-07 | 4.45 | 0.62 |
| Kcne3 | 2.41E-13 | 4.67 | 0.73 |
| Lsr | 3.94E-07 | 4.20 | 0.62 |
| Fam8a1 | 0.00023 | 3.99 | 0.48 |
| Polr3h | 0.00250 | 3.95 | 0.41 |
| Ctsh | 0.00014 | 3.80 | 0.49 |
| Gper1 | 1.57E-07 | 3.57 | 0.63 |
| Bgn | 2.75E-06 | 3.47 | 0.58 |
| Efna5 | 1.97E-05 | 3.44 | 0.54 |
| Lmo1 | 0.00027 | 3.44 | 0.48 |
| Eln | 0.00016 | 3.87 | 0.76 |
| Sdc1 | 6.44E-06 | 3.34 | 0.47 |
| Vegfc | 0.00026 | 3.33 | 0.57 |
| Clu | 0.00046 | 3.28 | 0.54 |
| Serpinf1 | 1.01E-06 | 3.18 | 0.68 |
| Dkk2 | 5.13E-05 | 3.47 | 0.80 |
| Cthrc1 | 2.41E-08 | 3.15 | 0.75 |
