## Supplemental Figures for "Developmental diversity and unique sensitivity to injury of lung endothelial subtypes during a period of rapid postnatal growth"

### Slide 1
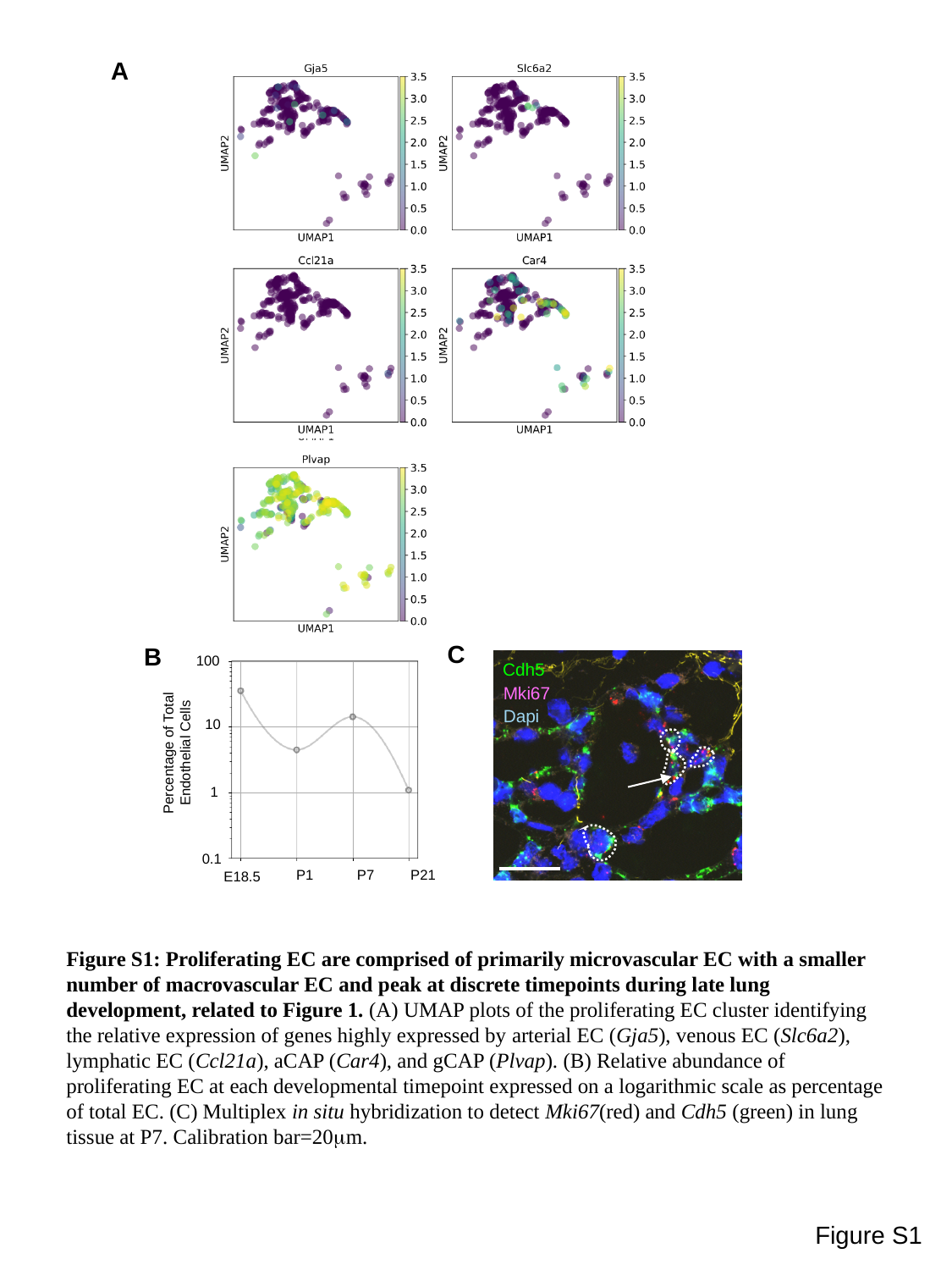

A
C
B
100
10
Percentage of Total Endothelial Cells
1
0.1
P1
P7
P21
E18.5
Cdh5
Mki67
Dapi
Figure S1: Proliferating EC are comprised of primarily microvascular EC with a smaller number of macrovascular EC and peak at discrete timepoints during late lung development, related to Figure 1. (A) UMAP plots of the proliferating EC cluster identifying the relative expression of genes highly expressed by arterial EC (Gja5), venous EC (Slc6a2), lymphatic EC (Ccl21a), aCAP (Car4), and gCAP (Plvap). (B) Relative abundance of proliferating EC at each developmental timepoint expressed on a logarithmic scale as percentage of total EC. (C) Multiplex in situ hybridization to detect Mki67(red) and Cdh5 (green) in lung tissue at P7. Calibration bar=20μm.
Figure S1

### Slide 2
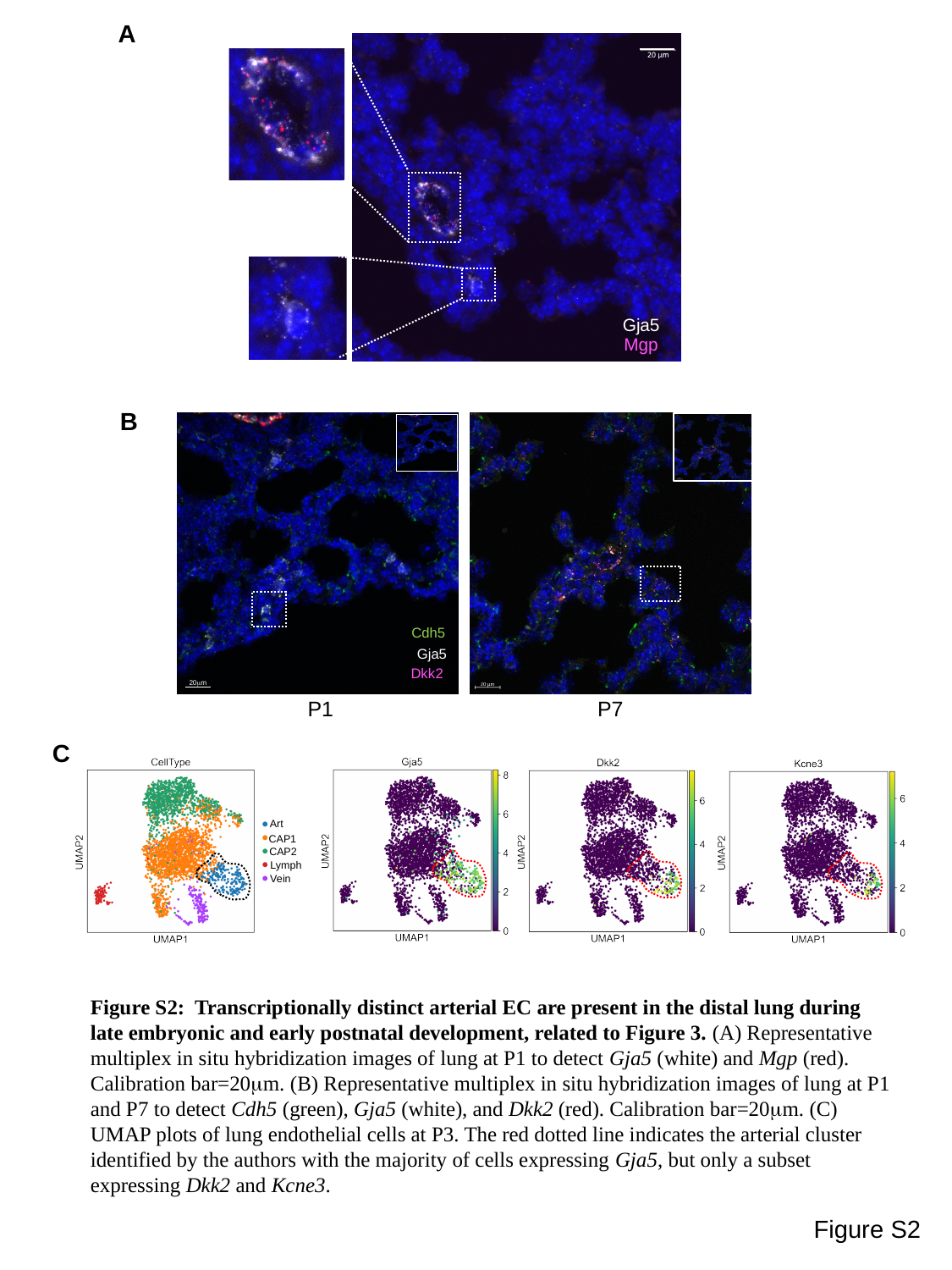

A
Gja5
Mgp
B
20μm
P7
P1
Cdh5
Gja5
Dkk2
C
Art
CAP1
CAP2
Lymph
Vein
Figure S2: Transcriptionally distinct arterial EC are present in the distal lung during late embryonic and early postnatal development, related to Figure 3. (A) Representative multiplex in situ hybridization images of lung at P1 to detect Gja5 (white) and Mgp (red). Calibration bar=20μm. (B) Representative multiplex in situ hybridization images of lung at P1 and P7 to detect Cdh5 (green), Gja5 (white), and Dkk2 (red). Calibration bar=20μm. (C) UMAP plots of lung endothelial cells at P3. The red dotted line indicates the arterial cluster identified by the authors with the majority of cells expressing Gja5, but only a subset expressing Dkk2 and Kcne3.
Figure S2

### Slide 3
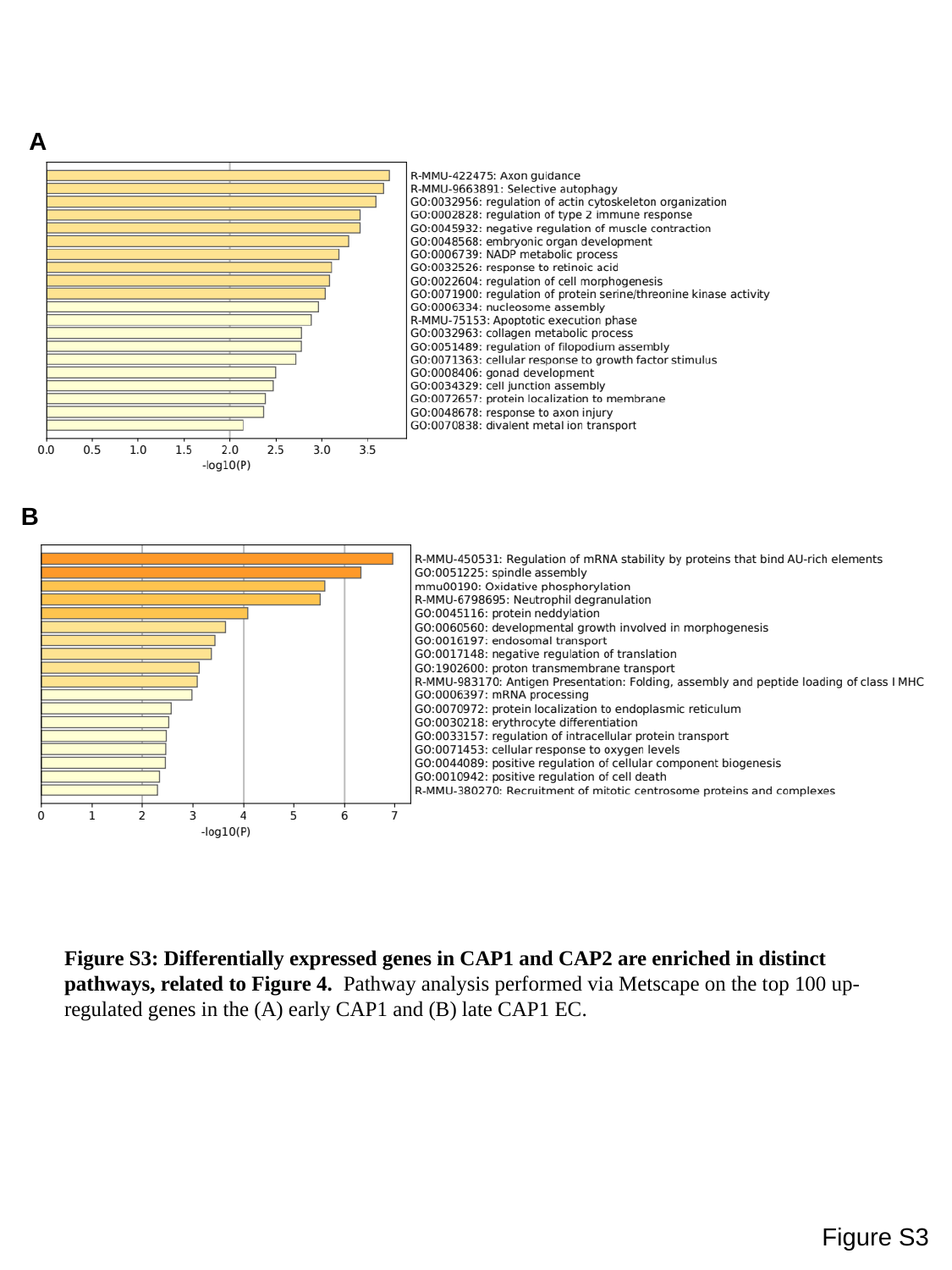

A
B
Figure S3: Differentially expressed genes in CAP1 and CAP2 are enriched in distinct pathways, related to Figure 4. Pathway analysis performed via Metscape on the top 100 up-regulated genes in the (A) early CAP1 and (B) late CAP1 EC.
Figure S3

### Slide 4
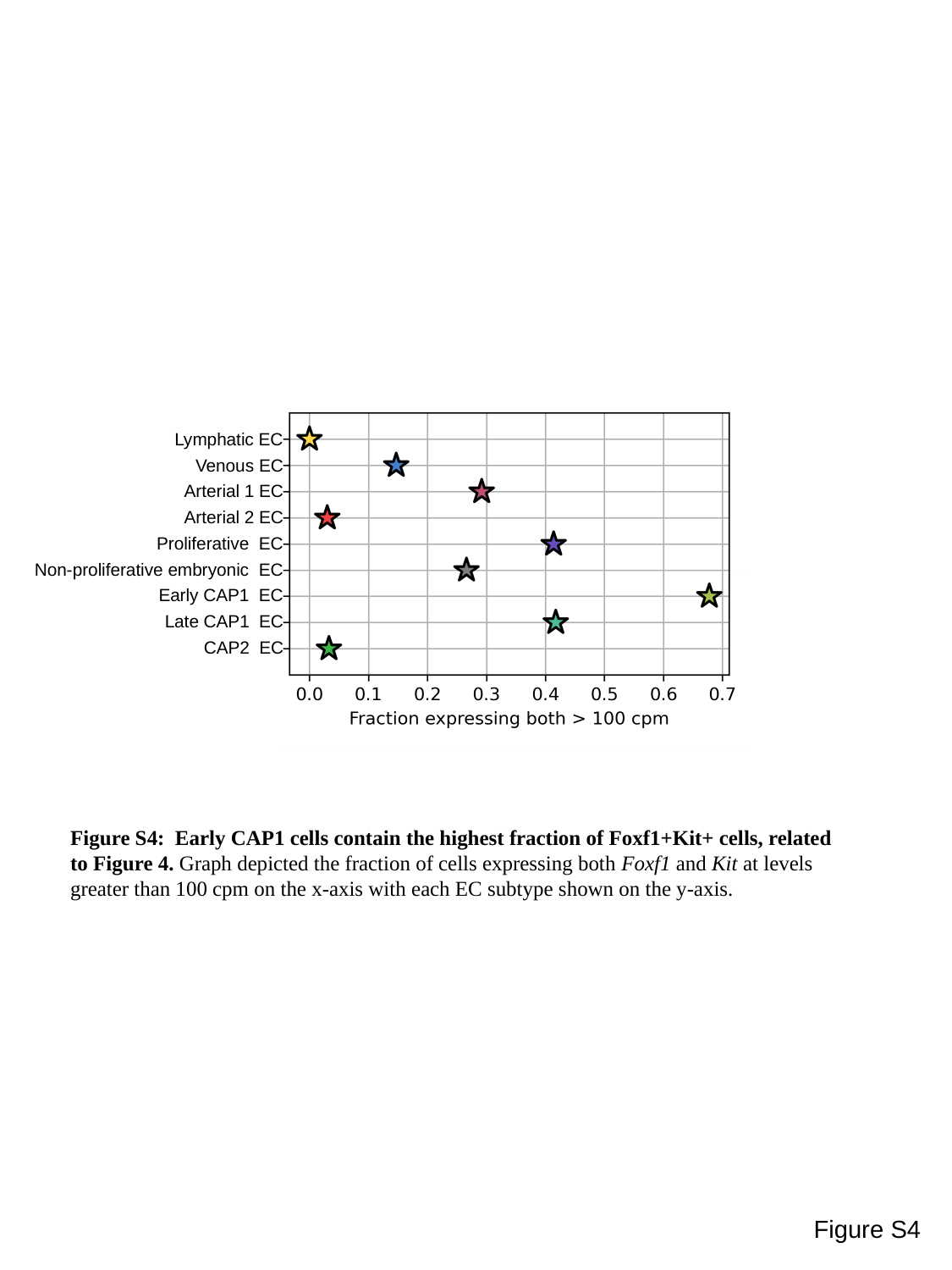

Lymphatic EC
Venous EC
Arterial 1 EC
Arterial 2 EC
Proliferative EC
Non-proliferative embryonic EC
Early CAP1 EC
Late CAP1 EC
CAP2 EC
Figure S4: Early CAP1 cells contain the highest fraction of Foxf1+Kit+ cells, related to Figure 4. Graph depicted the fraction of cells expressing both Foxf1 and Kit at levels greater than 100 cpm on the x-axis with each EC subtype shown on the y-axis.
Figure S4

### Slide 5
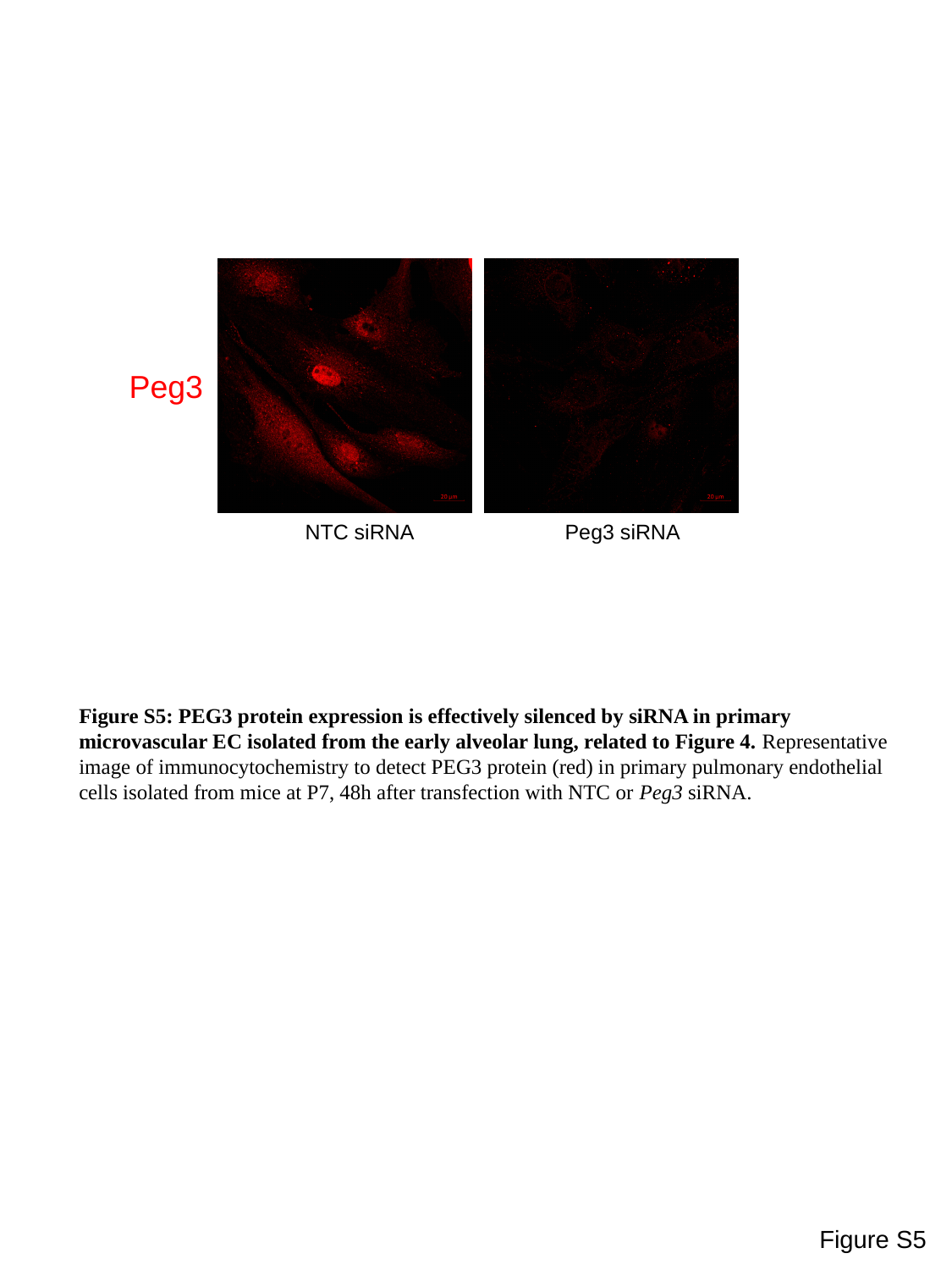

Peg3
NTC siRNA
Peg3 siRNA
Figure S5: PEG3 protein expression is effectively silenced by siRNA in primary microvascular EC isolated from the early alveolar lung, related to Figure 4. Representative image of immunocytochemistry to detect PEG3 protein (red) in primary pulmonary endothelial cells isolated from mice at P7, 48h after transfection with NTC or Peg3 siRNA.
Figure S5

### Slide 6
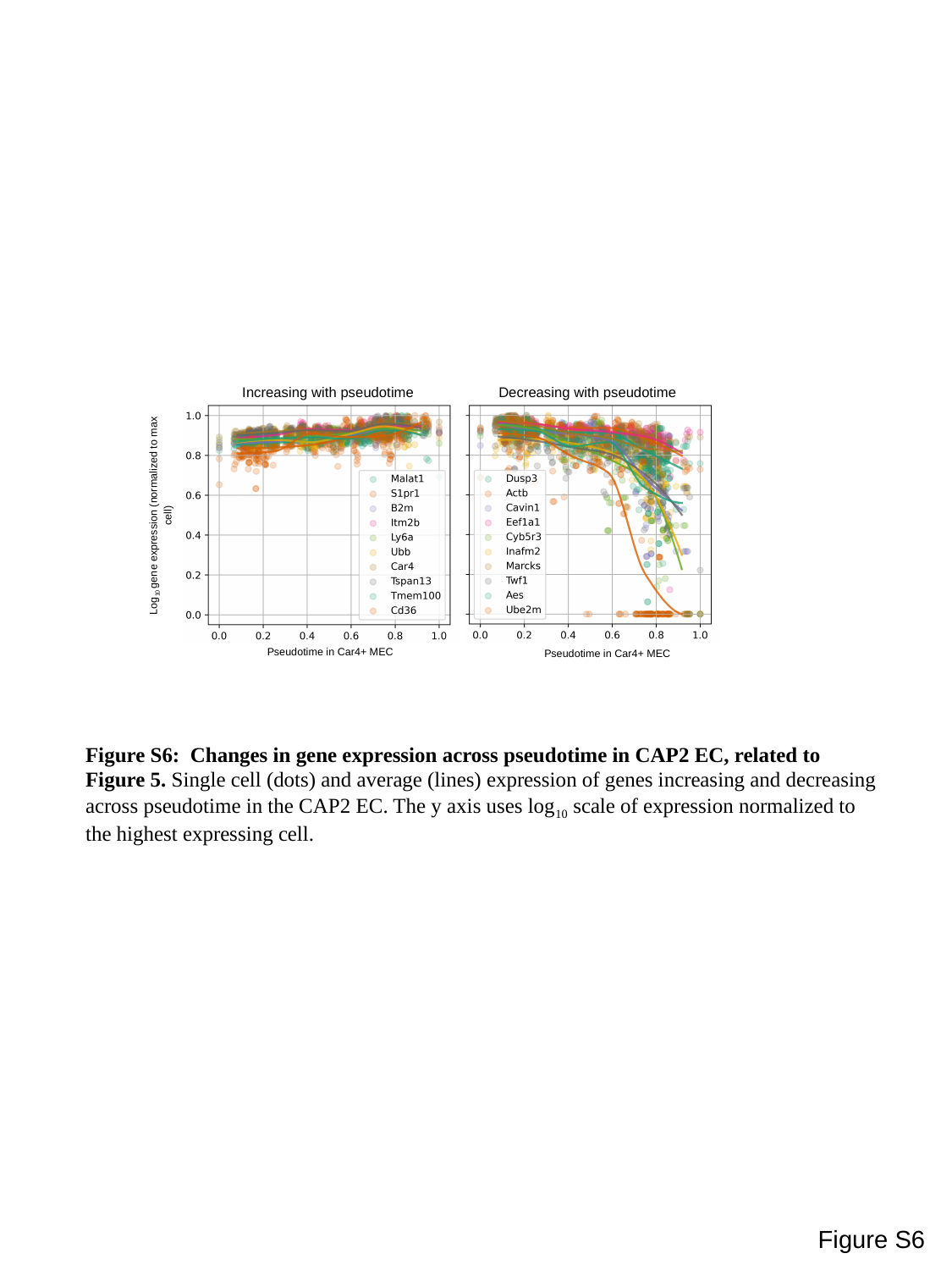

Increasing with pseudotime
Decreasing with pseudotime
Log_10_ gene expression (normalized to max cell)
Pseudotime in Car4+ MEC
Pseudotime in Car4+ MEC
Figure S6: Changes in gene expression across pseudotime in CAP2 EC, related to Figure 5. Single cell (dots) and average (lines) expression of genes increasing and decreasing across pseudotime in the CAP2 EC. The y axis uses log_10_ scale of expression normalized to the highest expressing cell.
Figure S6

### Slide 7
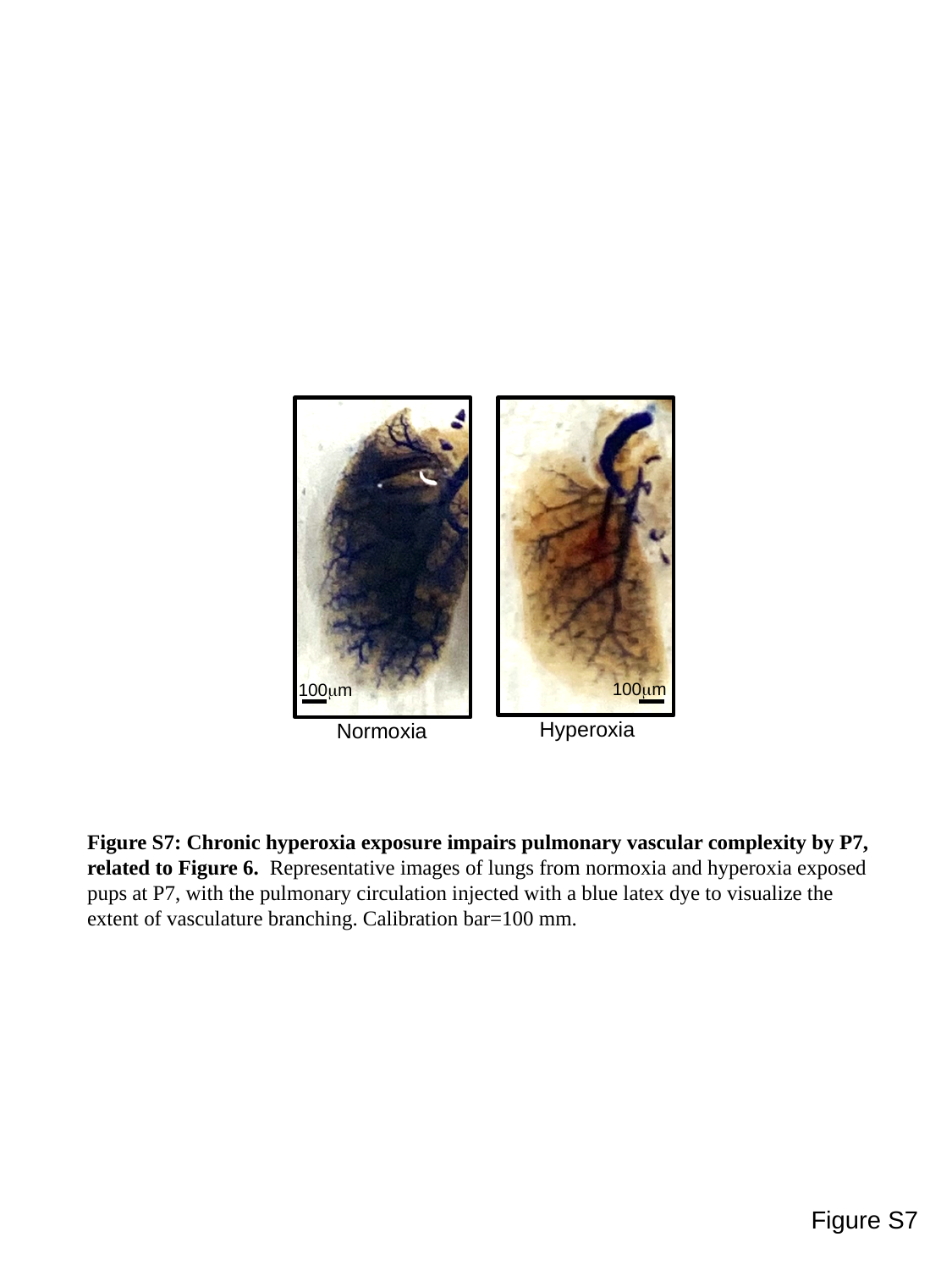

100μm
100μm
Hyperoxia
Normoxia
Figure S7: Chronic hyperoxia exposure impairs pulmonary vascular complexity by P7, related to Figure 6. Representative images of lungs from normoxia and hyperoxia exposed pups at P7, with the pulmonary circulation injected with a blue latex dye to visualize the extent of vasculature branching. Calibration bar=100 mm.
Figure S7

### Slide 8
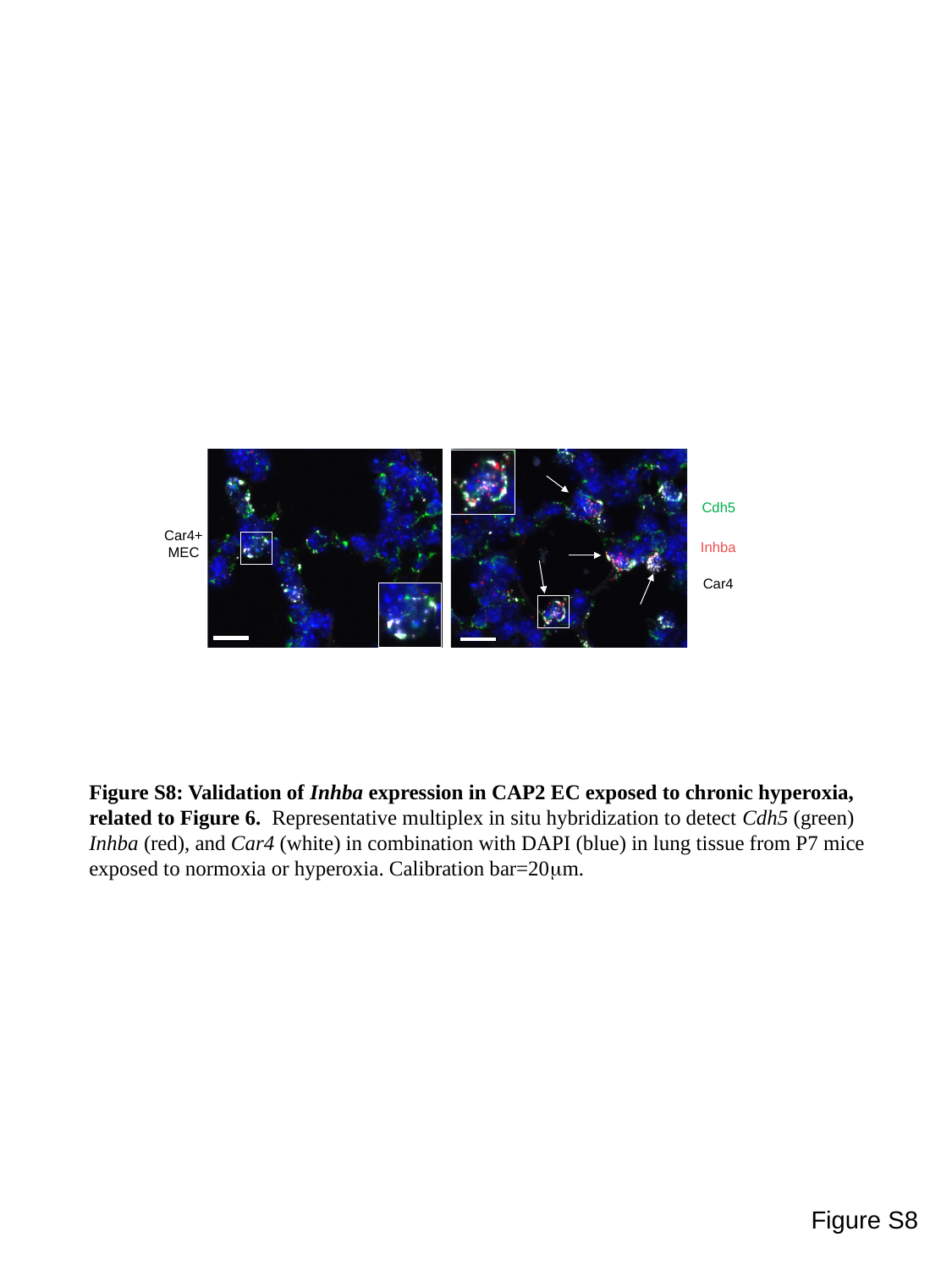

Cdh5
Car4+MEC
Inhba
Car4
Figure S8: Validation of Inhba expression in CAP2 EC exposed to chronic hyperoxia, related to Figure 6. Representative multiplex in situ hybridization to detect Cdh5 (green) Inhba (red), and Car4 (white) in combination with DAPI (blue) in lung tissue from P7 mice exposed to normoxia or hyperoxia. Calibration bar=20μm.
Figure S8
